# Subcellular localization of GPCR kinases differentially modulate biased signaling at CXCR3

**DOI:** 10.1101/2022.07.11.499601

**Authors:** Julia Gardner, Dylan Scott Eiger, Chloe Hicks, Issac Choi, Uyen Pham, Sudarshan Rajagopal

## Abstract

Some G protein-coupled receptors (GPCRs) demonstrate *biased signaling*, where ligands of the same receptor differentially activate specific downstream signaling pathways over others. Ligand-specific receptor phosphorylation by GPCR kinases (GRKs) is one mechanism underlying this phenomenon. Recent evidence demonstrates that GPCRs traffic to and signal from subcellular compartments beyond the plasma membrane, a paradigm termed *location bias*. Here, we show that GRKs translocate to endosomes following stimulation of the chemokine receptor CXCR3 and other GPCRs. The GRK recruitment patterns at the plasma membrane and endosome are distinct and depend on the identity of the ligand used to activate the receptor. Using cells deficient of GRKs, we demonstrate that biased ligands have unique signaling profiles upon rescue of location-specific GRK isoforms. Our work highlights a role of the GRKs in location-biased GPCR signaling and demonstrates the complex interactions between ligand, GRK isoform and cellular location that contribute to biased signaling.

---

G protein-coupled receptors (GPCRs) represent the largest and most versatile class of receptors in humans and can elicit cellular responses to various stimuli, including from photons, odorants, peptides, lipids, and small molecules^1^. Due to the extensive involvement of GPCRs in normal physiology and pathophysiology, approximately 35% of all Food and Drug Administration approved drugs target one or more of the 800 human GPCRs^2, 3^. Although GPCRs engage an array of signaling effectors to actuate a cellular response, such as proteins, GPCR kinases (GRKs), and G β-arrestins, the mechanisms underlying GPCR signaling are still incompletely understood^4^.

GPCRs are conventionally studied at the plasma membrane, where ligand binding to an orthosteric pocket promotes a conformational change in the receptor that induces intracellular activation of G proteins and subsequent G protein-dependent signaling^5, 6^. GRKs and other kinases then phosphorylate the receptor C-terminus and intracellular loops, promoting the binding of β-arrestins, which desensitize receptor coupling to G proteins and induce receptor internalization into clathrin-coated pits^7, 8^. β-arrestins can also act as scaffolds for other signaling effectors, including mitogen activated protein kinases (MAPKs) and E3 ubiquitin ligases^9, 10^.

Evidence shows that some ligand:GPCR complexes can preferentially activate distinct signaling pathways over others, a phenomenon referred to as *biased signaling*. Biased signaling can be driven by the ligand, receptor, or tissue/cell (system)^11–13^. One proposed mechanism for biased signaling, the *phosphorylation barcode hypothesis*, posits that unique ligand-induced patterns of GPCR phosphorylation promote selective engagement with specific signaling effectors, which produce functionally selective outputs^14–16^. Recent studies demonstrate the critical role the phosphorylation barcode plays in driving specific β-arrestin recruitment and conformational signatures, ultimately promoting unique downstream signaling^17–20^.

Surprisingly, despite the known function of GRKs in receptor phosphorylation, the role of the GRKs in promoting biased signaling is poorly understood. Experimental attempts to study the GRKs are obscured by the complexity of the protein system itself. The seven identified mammalian GRKs (1-7) are divided into three subfamilies according to gene sequence and structure^21^, wherein GRKs 1 and 7 comprise the GRK1 family, GRKs 2 and 3 the GRK2 family, and GRKs 4, 5, and 6 the GRK4 family^22, 23^. GPCRs demonstrate immense heterogeneity in the putative phosphorylation patterns present on intracellular loop 3 (ICL3) and C-termini^24^, and explication of how these seven GRKs interact with hundreds of GPCRs remains elusive^25^. GRKs 2, 3, 5, and 6 are ubiquitously expressed in human tissues^26^, and while no specific consensus sequence has been identified for specific GRK isoforms^27^, recent work has demonstrated that GRKs 2, 3, 5 and 6 each possess unique signaling roles at GPCRs and other substrate proteins including non-GPCR receptors^28^. Interestingly, GRKs 2 and 3 are expressed cytosolically, and ligand-induced engagement with a GPCR is dependent upon interaction of the GRK pleckstrin homology (PH) domain with free Gβγ subunit while GRKs 5 and 6 are constitutively expressed on the plasma membrane^30, 31^.

It has been difficult to experimentally delineate the specific contributions of individual GRKs to the generation of unique GPCR phosphorylation patterns and subsequent receptor regulation. Studies using phosphosite-specific antibodies or mass spectrometry have identified certain residues on numerous GPCRs that are specifically targeted by individual GRKs^15, 32, 33^. However, other sites have been shown to be phosphorylated by multiple GRKs^34^, and it remains unknown if the order in which a receptor engages GRK isoforms modulates the ultimate phosphorylation pattern. Differential GRK expression levels in various tissues have also been proposed as a means of attaining GRK-specific functionality. For example, GRK5 and GRK6 are expressed commensurately in B- and T-cells, yet GRK5 is the predominantly expressed isoform in cardiac tissue^27^. While distinct structural elements, phospho-site targets, and tissue expression levels of the GRKs have helped understand isoform specificity, further studies are needed to fully resolve this complex system.

It has been recently demonstrated that GPCRs can signal from numerous subcellular locations beyond the plasma membrane, including the endosome, mitochondria, Golgi apparatus, nuclear membrane, and endoplasmic reticulum^35–41^. For example, we previously demonstrated that biased endogenous ligands of the chemokine receptor CXCR3 induce distinct patterns of G protein activation and β-arrestin recruitment at endosomes compared to the plasma membrane, and CXCR3 internalization contributes to the overall biased cellular output at this receptor^42^. However, it is incompletely understood how GPCRs elicit unique signaling events in a location-dependent manner. Given the critical role of receptor phosphorylation in biased GPCR signaling and initiating receptor internalization cascades, it is likely that the GRKs play a substantial role in generating functionally selective responses in different cellular locations. To our knowledge, few experiments have assessed if the GRKs can engage with GPCRs beyond the plasma membrane.

Here, we show that biased CXCR3 small-molecule agonists promote different patterns of GRK engagement with CXCR3 at the plasma membrane. We also provide evidence demonstrating that some ligands can promote GRK translocation to endosomes in a pattern entirely distinct from that observed at the plasma membrane. We further show that individual GRKs exhibit distinct effects on CXCR3 signaling depending on the biased ligands used to activate the receptor. We show association of GRKs 2 and 3 with CXCR3 is largely dependent upon activation of G proteins, but some ligands enable G protein-independent mechanisms of recruitment. Each GRK demonstrates unique roles in regulating the receptor’s ability to engage β-arrestins, promote receptor internalization, and activate extracellular-signal-regulated kinase (ERK). Using engineered GRK-location mutants, we show that GRK identity and subcellular localization modulate GPCR signaling depending on the ligand used to activate the receptor. Lastly, we demonstrate that GRK recruitment to endosomes is not unique to CXCR3 but is observed across a wide range of GPCRs, suggesting that location-specific GRK activity is observed across the GPCR superfamily. These findings suggest that subcellular engagement of GRKs with GPCRs may facilitate location-specific signaling responses, and that the role of individual GRKs are differentially directed by different ligands. These data suggest a complex interaction between GRK isoforms, ligands, and cellular locations in driving a GPCR’s overall biased signaling outputs.

## RESULTS

### Biased ligands demonstrate different GRK recruitment patterns to the plasma membrane and CXCR3

CXCR3 is a chemokine receptor with three known endogenous ligands that exhibit biased signaling in various forms, such as in their differential formation of G⍰i: β-arrestin complexes and markedly different abilities to induce G protein- or β-arrestin-mediated signaling^43–46^. We first determined if activation of CXCR3 using biased ligands promotes distinct patterns of GRK recruitment to the plasma membrane in HEK293 cells. Using a previously validated nanoBiT complementation assay^47^, we monitored the interaction between a plasma membrane marker CD8⍰-SmBiT and GRK2-, GRK3-, GRK5-, or GRK6-LgBiT following stimulation of untagged CXCR3 (**Figure 1A**). We used three endogenous, biased CXCR3 agonists, CXCL9, CXC10, and CXCL11, and two synthetic CXCR3 agonists, VUF10661 and VUF11418 for these experiments^48^. CXCL11 and VUF10661 are relatively β-arrestin-biased while CXCL10 and VUF11418 are relatively G protein-biased. CXCL9 acts as a partial β-arrestin-biased agonist^49^.

**Figure 1:**
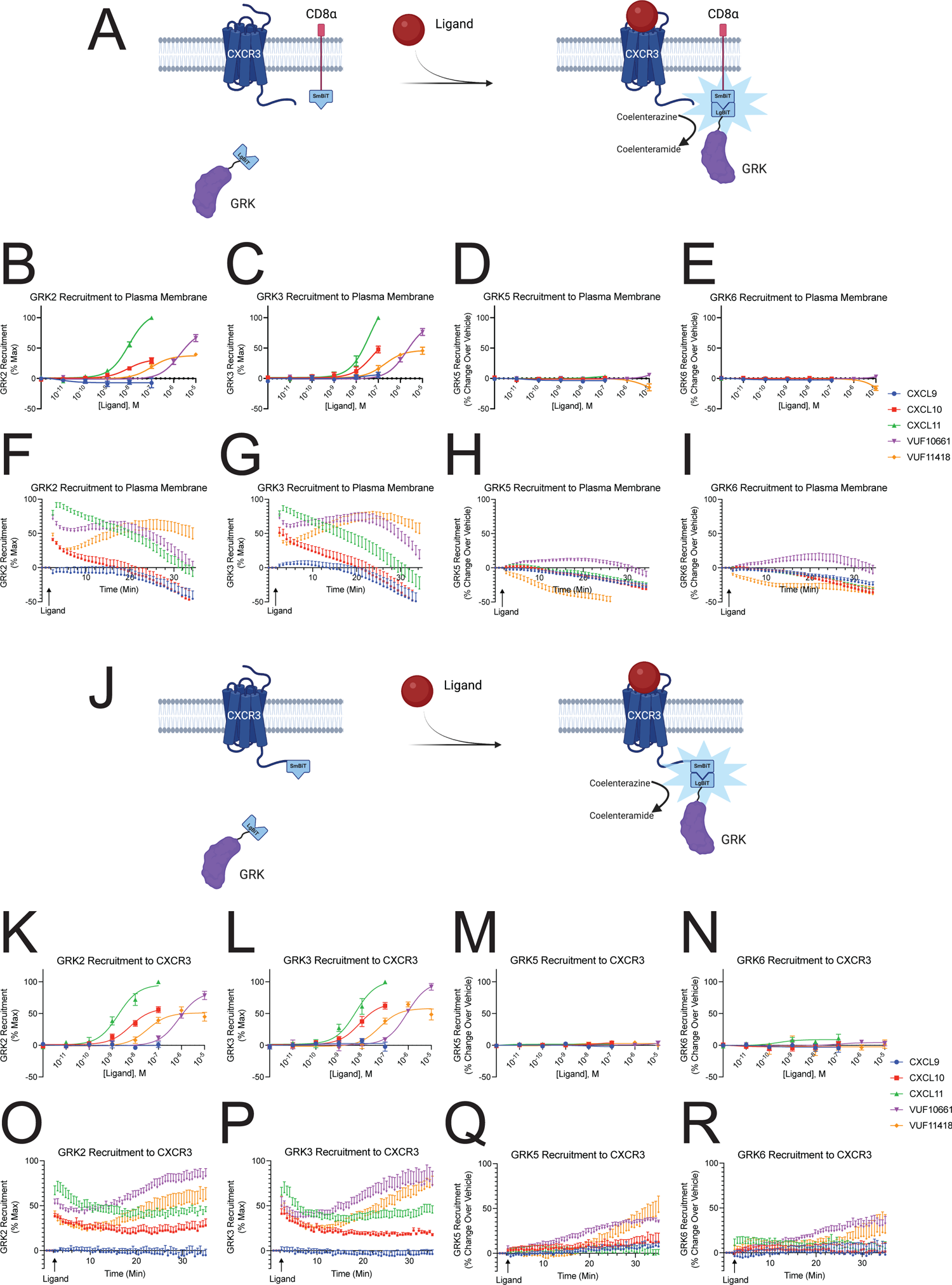
GRK recruitment to the plasma membrane and CXCR3. **(A)** Schematic representation of a NanoBiT complementation assay detecting GRK recruitment to the plasma membrane using GRK-LgBiT and CD8⍰-SmBiT in HEK293 cells. **(B-E)** Agonist concentration-response and **(F-I)** kinetic time-course of GRK2, GRK3, GRK5, and GRK6 recruitment to the plasma membrane. **(J)** Schematic representation of a NanoBiT complementation assay detecting GRK recruitment to CXCR3 using GRK-LgBiT and CXCR3-SmBiT in HEK293 cells. (**K to N**) Agonist concentration-response and (**O to R**) kinetic time-course of GRK2, GRK3, GRK5, and GRK6 recruitment to CXCR3. All experiments were performed following stimulation with either 100nM CXCL9, CXCL10, or CXCL11, and 10µM VUF10661 or VUF11418. Data shown are the mean ± SEM of n=5. Concentration-response curves for GRK2 and GRK3 are normalized to maximum signal while concentration-response curves for GRK5 and GRK6 are shown as luminescence change over vehicle.

After 5 minutes, the recruitment patterns of GRK2 and GRK3 were similar to each other, with differences in efficacy and potency between the biased ligands (**Figure 1B and 1C**). Specifically, CXCL11 and VUF10661 were the most efficacious agonists followed by CXCL10 and VUF11418. CXCL9 demonstrated little to no detectable recruitment of GRK2 or GRK3 at physiologic doses of chemokine. GRK2/3 and GRK5/6 are known to localize differentially in the cell at baseline, so we performed confocal microscopy using YFP-tagged GRK2, GRK3, GRK5, or GRK6 and CXCR3-mCerulean to confirm cytosolic expression of GRK2 and GRK3 and constitutive membrane localization of GRK5 and GRK6^50–52^ (**Supplemental Figure 1**). Thus, ligand-induced recruitment to the plasma membrane is not necessarily an illustrative readout of GRK5 and GRK6 activity, as evinced by the lack of detectable recruitment of these GRKs at physiological chemokine doses in our assay (**Figure 1D and 1E**).

In analyzing the kinetic data of GRK recruitment to the plasma membrane, we observed significant differences across ligands. All ligands, except CXCL9, demonstrate rapid recruitment of GRK2 and GRK3 to the plasma membrane within seconds of receptor activation (**Figure 1F-1I**). However, at 30 minutes, all ligands except VUF11418, show a marked decrease in luminescence. These data demonstrate that the ligands not only can promote different magnitudes of GRK2 and GRK3 translocation upon receptor activation, but also can produce unique kinetic signatures for the interaction of individual GRKs with the plasma membrane. These data suggest that differential GRK recruitment magnitude and kinetics may contribute to the generation of specific phosphorylation barcodes and biased signaling.

When the same experiment was performed using CXCR3-SmBiT to assess GRK recruitment to the receptor, we observed nearly identical dose response curves for all GRKs 5 minutes after stimulation (**Figure 1J-1R**). Surprisingly when analyzing GRK recruitment over time, we observed an increase in luminescence signal at 30 minutes for all four GRKs using VUF10661 and VUF11418 but not the endogenous chemokines (**Figure 1O-1R**). For GRK2 and GRK3, following an initial robust phase of GRK recruitment induced by these synthetic ligands, there was a slight decrease in luminescence at 10 minutes, followed by an increase in luminescence at 30 minutes (**Figure 1O-1P**). CXCL10 and CXCL11 demonstrated sustained interaction with the receptor even at 30 minutes – these data differed when measuring GRK recruitment to the plasma membrane where the luminescent signal returned to baseline at this time. By contrast, GRK5 and GRK6 showed little initial activity following stimulation with these small-molecules but demonstrate a very slow and late recruitment phase after 10 minutes (**Figure 1Q-1R**).

### GRK recruitment patterns to endosomes differs than those observed at the plasma membrane

Based on the unexpected kinetic tracings of GRK recruitment to CXCR3 compared to the plasma membrane, we hypothesized this data may represent GRKs interacting with the receptor at subcellular locations. To evaluate potential recruitment of the GRKs to endosomes, we repeated the NanoBiT complementation assay using the indicated GRK-LgBiT, WT CXCR3, and the previously validated early endosome marker 2x-Fyve tagged with SmBiT^53, 54^ (**Figure 2A**). We observed that GRK2, GRK3, and GRK5 could be recruited to endosomes, with VUF10661 exhibiting the most robust recruitment of these GRKs (**Figure 2B-2E**). We also observed detectable recruitment of GRK3 to endosomes using CXCL11, VUF11418, and CXCL10. The overall pattern of ligand-induced GRK recruitment to endosomes significantly differed from the recruitment pattern observed at the plasma membrane. GRK recruitment to endosomes was significantly slower, consistent with previously reported kinetic data on GPCR internalization^55, 56^.

**Figure 2:**
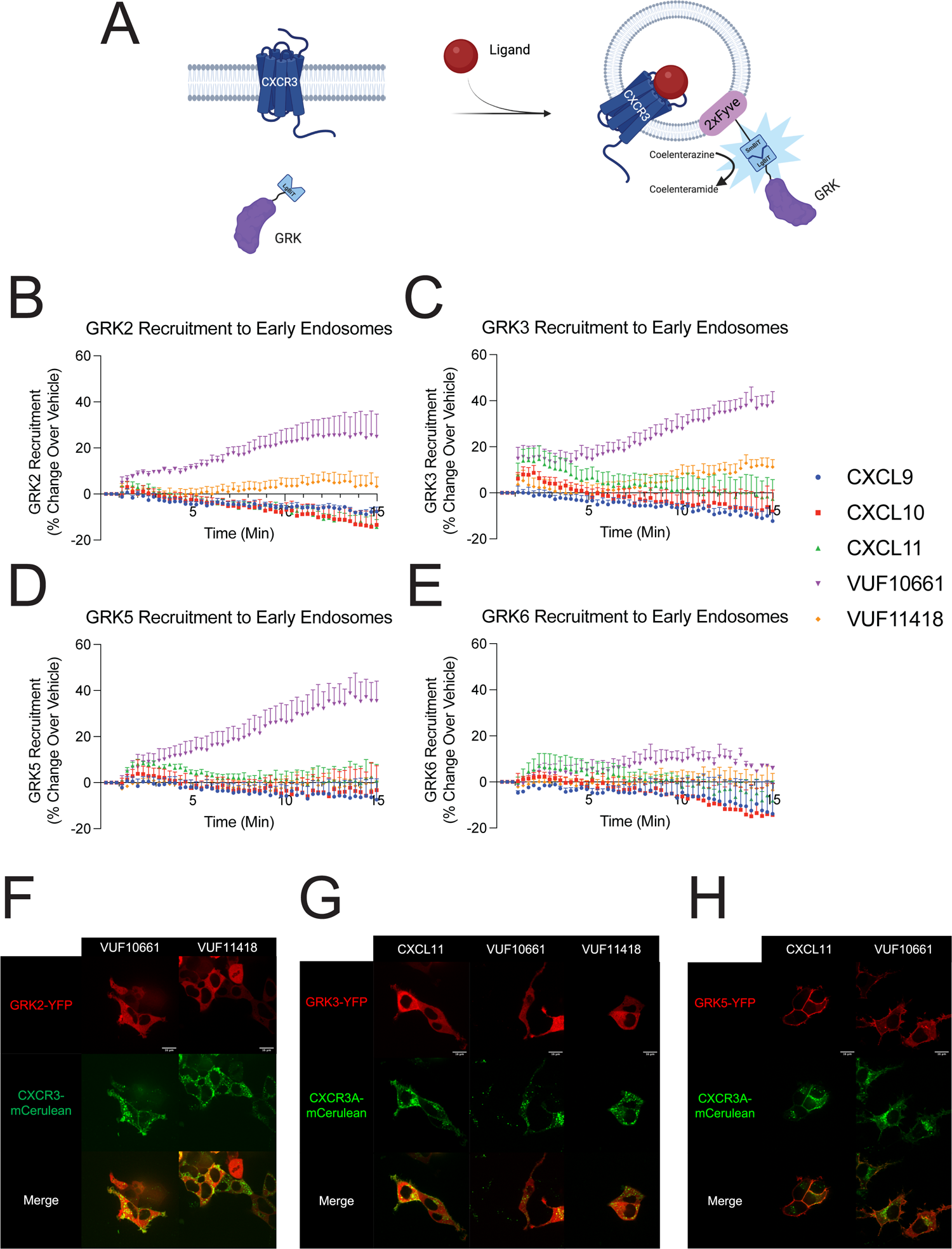
CXCR3-mediated GRK recruitment to the endosome and the plasma membrane. **(A)** Schematic representation of the NanoBiT complementation assay used to detect GRK recruitment to endosomes using wild-type CXCR3, LgBiT-tagged GRK, and 2xFyve-SmBiT in HEK293 cells. Following activation of the receptor and recruitment of GRK to endosomes, the smBiT and LgBiT undergo complementation and generate a functional luminescent signal as a readout of GRK recruitment to endosome. Kinetic data of (**B**) GRK2, (**C**) GRK3, (**D**) GRK5, and (**E**) GRK6 recruitment levels to early endosomes upon stimulation with either 100nM CXCL9, CXCL10, and CXCL11, and 10µM VUF10661 or VUF11418. Kinetic tracings are shown as luminescence change over vehicle. **(F to H)** Confocal microscopy images of HEK293 cells transfected with CXCR3-mCerulean and (**F**) GRK2-YFP, (**G**) GRK3-YFP, or (**H**) GRK5-YFP. Images were taken 45 minutes after stimulation with 100nM of CXCL11 or 10µM of VUF10661 or 10µM of VUF11418. Plate-based experiments show the mean ± SEM of n=5. Confocal microscopy images are representative of n=3.

To validate these findings with confocal microscopy, we transfected HEK293 cells with the indicated GRK-YFP (shown in red) and CXCR3-mCerulean (shown in green) and stimulated with agonist for 45 minutes (**Figure 2F-2H**). While all ligands did promote receptor internalization, we observed the formation of intracellular yellow puncta, suggestive of colocalized receptor and GRK in endosomes, only for some ligand:GRK combinations, consistent with our luminescence data **(Supplemental Figures 2 and 3**). These data provide evidence that CXCR3 can engage GRKs in endosomes. Of note, we observed colocalization of GRK3-YFP and CXCR3-mCerulean following treatment with CXCL11 but not with CXCL10 nor CXCL9 (**Supplemental Figure 2B**). In contrast, following treatment with VUF10661, we observed robust colocalization of CXCR3-mCerulean and GRK2-YFP, GRK3-YFP, and to a lesser extent GRK5-YFP. These findings suggest that ligands may direct isoform-specific activities of the GRKs through selective trafficking of certain isoforms to endosomes.

**Figure 3:**
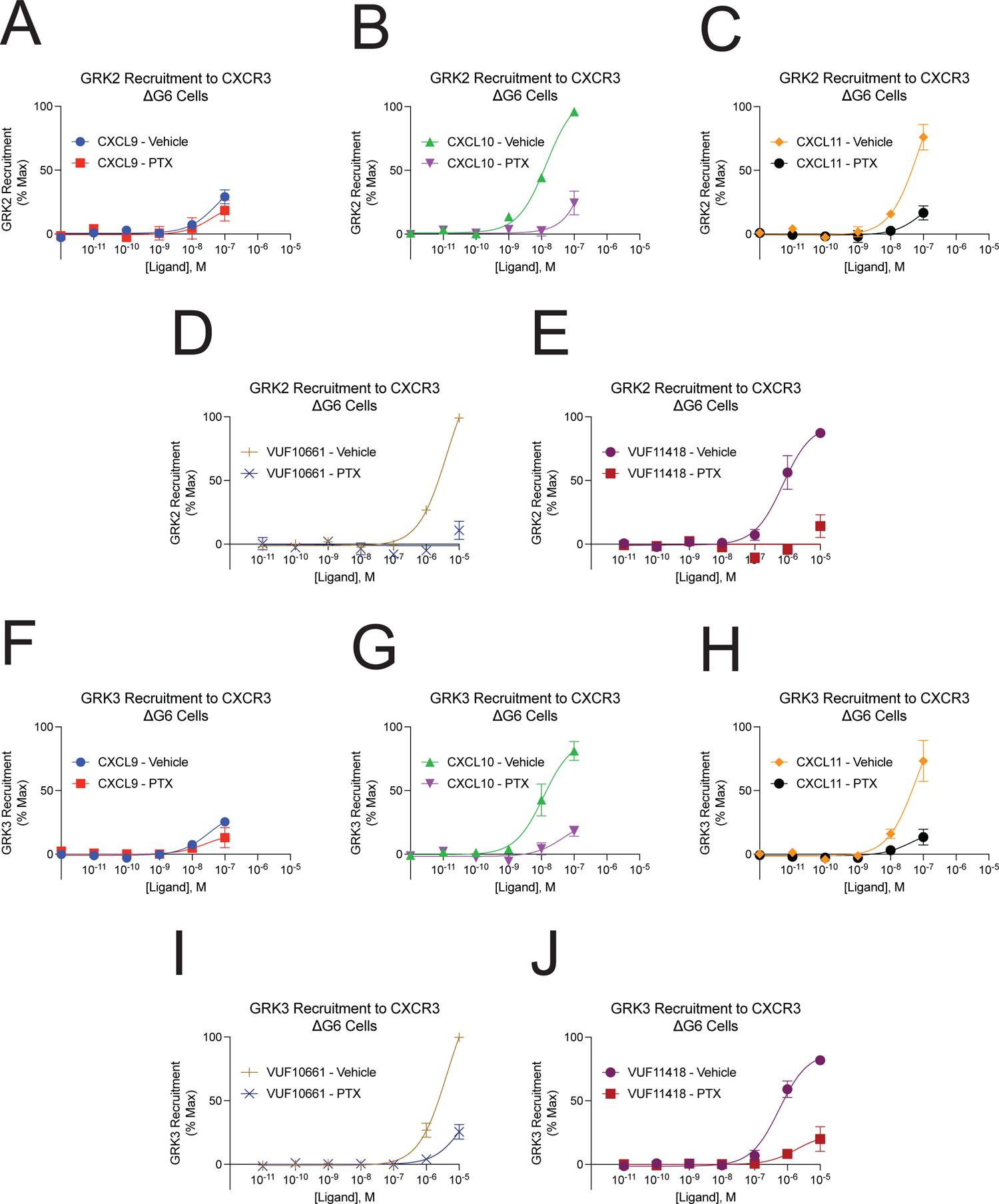
GRK recruitment to CXCR3 in G protein-deficient cells (ΔG6 cells). Agonist concentration-response of **(A-E)** GRK2 and **(F-J)** GRK3 recruitment to CXCR3 with and without pretreatment with 200ng/mL pertussis toxin (PTX) in ΔG6 knockout cells treated with 100nM CXCL9, CXCL10, or CXCL11, and 10µM VUF10661 or VUF11418. ΔG6 is a HEK293 cell line devoid of six G proteins: *GNAS*, *GNAL*, *GNA11*, *GNA12*, *GNA13*, and *GNAQ*. GRK recruitment was measured using a split luciferase assay involving LgBiT-tagged GRK2 or GRK3, and CXCR3-SmBiT. Data shown are the mean ± SEM of n=5, *P<0.05 by two-way ANOVA with Tukey post hoc testing between treatment conditions at a single GRK:ligand combination. Concentration-response curves are normalized to maximum signal observed across all ligands.

### GRK2 and GRK3 recruitment to CXCR3 is largely dependent upon G protein activation, but some ligands demonstrate G protein-independent recruitment mechanisms

A primary feature distinguishing the GRK2 and GRK4 subfamilies is the reliance of GRK2 and GRK3 recruitment to a GPCR upon interactions of their PH domain with free Gβγ following heterotrimeric G protein dissociation^57, 58^. This model proposes that β-arrestin binding is tied to G protein-mediated GRK recruitment. Although studies have shown that mutations in the PH domain of GRK2 abolish its ability to interact with free Gβγ and phosphatidylinositol 4,5-bisphosphate^59^, various experiments have also suggested G protein-independent mechanisms of GRK recruitment. For example, the atypical chemokine receptor 3 (ACKR3), which does not activate G proteins, has been shown to recruit and activate GRKs 2, 3 and 5^60^. Further evidence suggests a role of direct, G protein-independent GRK2 recruitment to the D2R in facilitating β-arrestin engagement (Pack, et al., 2018). However, it is not yet clear if this effect is conserved across GPCRs or unique to a few receptor subtypes.

To evaluate the necessity of G protein activation in GRK2 and GRK3 recruitment to CXCR3, we assessed if GRKs could still be recruited to this predominantly G⍰i-coupled receptor in the absence of G⍰i ^61^. We took advantage of a previously validated HEK293 cell line (ΔG6 cells) devoid of all G proteins except G⍰i/o family members ^62^. Specifically, these cells lack *GNAS*, *GNAL*, *GNA11*, *GNA12*, *GNA13*, and *GNAQ*. Using these cells, we performed the NanoBiT complementation assay using GRK2- or GRK3-LgBiT and CXCR3- SmBiT with and without addition of pertussis toxin (PTX) to inhibit G⍰i/o activation^63, 64^ (**Figure 3A-3J**).

The kinetic tracings for GRK recruitment in the ΔG6 HEK293 cells were similar to that observed in WT HEK293 cells (**Supplemental Figure 4**). Surprisingly, we were able to detect GRK2 and GRK3 recruitment ΔG6 cells, a process we did not observe in WT HEK293 cells. We found that PTX treatment significantly reduces GRK2 and GRK3 recruitment, although there was incomplete inhibition of GRK2 recruitment using the endogenous chemokines (**Figure 3A-4C**), and GRK3 recruitment by all ligands (**Figure 3F-3J**). It is possible that these findings are due to incomplete inhibition of G⍰i/o activation with PTX treatment. However, we observed near complete inhibition of GRK2 recruitment with VUF10661 and VUF11418 (**Figure 3D and 3E**), but relatively little change with CXCL9 (**Figure 3A**), suggesting that PTX treatment was sufficient. These data validate previous reports suggesting that recruitment of GRK2 family proteins is largely, but not completely, dependent on G protein activation, with a component of direct, G protein-independent mechanisms for GRK2/3 recruitment.

**Figure 4:**
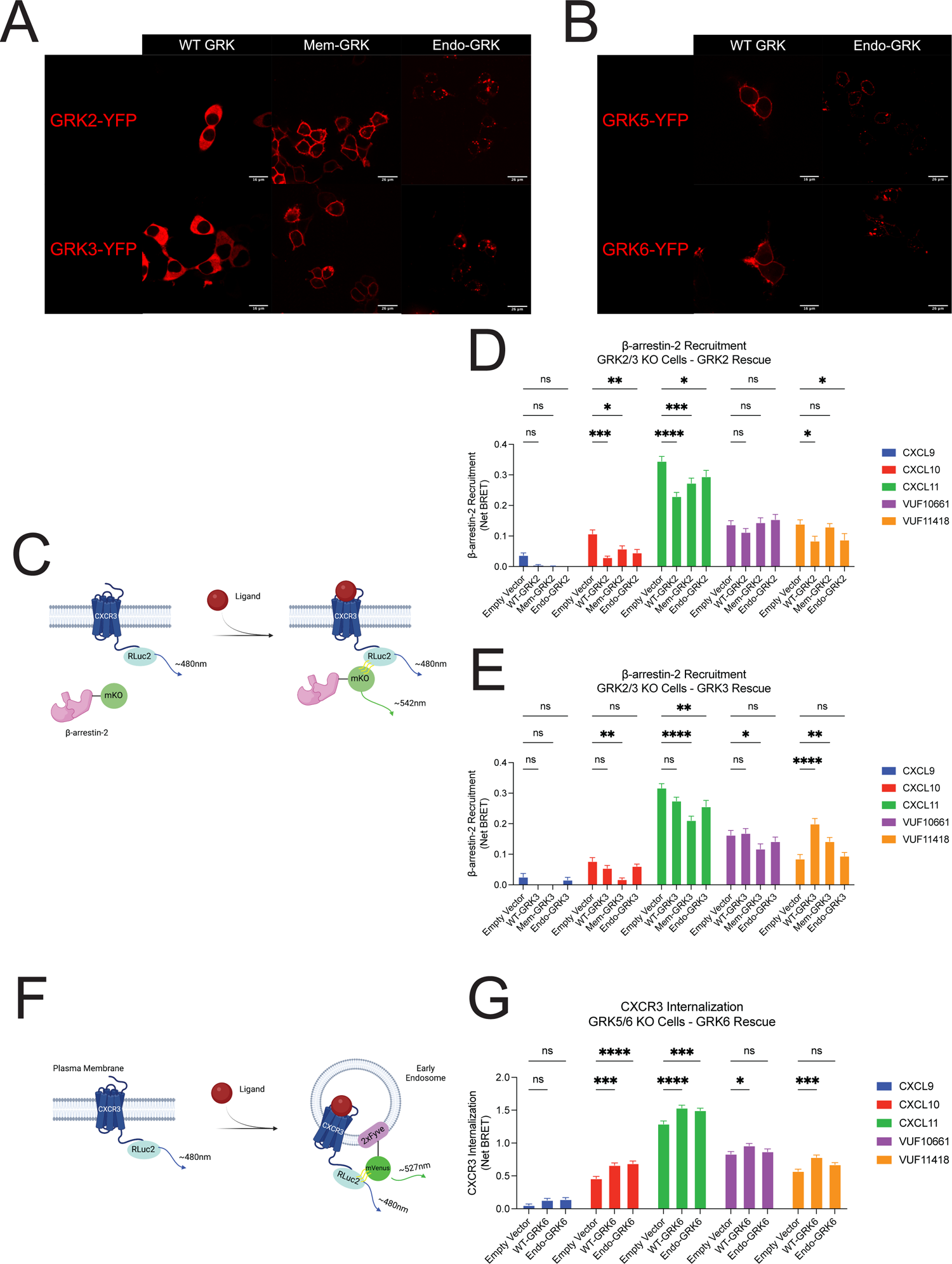
β-arrestin recruitment and CXCR3 receptor internalization in cells devoid of GRKs. Confocal microscopy images of HEK293T cells transfected with location-specific mutants of **(A)** GRK2 and GRK3, and **(B)** GRK5 and GRK6 tagged with a YFP (false red coloring for clarity). **(C)** Schematic representation of β-arrestin-2-mKO recruitment to CXCR3-Rluc2 using bioluminescence resonance energy transfer (BRET). BRET ΔGRK2/3 knockout cell line upon addback of **(D)** GRK2 and **(E)** GRK3 constructs or empty vector. **(F)** Schematic representation of BRET-based assay to assess CXCR3 internalization in ΔGRK5/6 upon addback of **(G)** GRK6 constructs or empty vector. 100nM CXCL9, CXCL10, or CXCL11, and 10µM VUF10661 or VUF11418 were used in all experiments. Data shown are the mean AUC ± SEM of n=5., *P<0.05 by two-way ANOVA with Tukey post hoc testing conducted between empty vector and other transfections conditions within each ligand.

To further corroborate these findings, we repeated these experiments in HEK293 cell line devoid of all G proteins (ΔG7 cells), specifically *GNAS*, *GNAL*, *GNAQ*, *GNA11*, *GNA12*, *GNA13*, *GNAI1*, *GNAI2*, *GNAI3*, *GNAO1*, *GNAZ*, *GNAT1*, and *GNAT2*^65^. Upon rescued expression of G⍰i1, we observed a significant increase in GRK2 and GRK3 recruitment at all ligands except for CXCL9 (**Supplemental Figure 5**). Even in the absence of G⍰i, VUF10661 and CXCL11 maintained a partial ability to recruit GRK2, while CXCL10, CXCL11, VUF10661, and VUF11418 all partially recruited GRK3. Together, these findings suggest an additional element of GRK isoform specificity, wherein different ligands can induce different mechanisms to promote GRK recruitment.

### GRK location-specific mutants differentially affect β-arrestin 2 recruitment and CXCR3 internalization

Given the findings that biased CXCR3 agonists promote differential GRK recruitment across subcellular locations, we next determined if the cellular localization of a GRK can influence receptor signaling. We generated location-specific GRK constructs to localize GRKs to either the plasma membrane or endosome. We used a membrane targeting sequence from the tyrosine protein kinase Lyn to create constitutively membrane-bound mutants (Mem-GRK) of GRK2 and GRK3, and a 2x-Fyve early endosome tag to develop endosome-bound mutants (Endo-GRK) of GRK2, GRK3, GRK5, and GRK6. Constructs were validated through confocal microscopy and showed plasma membrane localization of the Mem-GRK2 and Mem-GRK3 constructs, and endosome association of the four Endo-GRKs (**Figure 4A and 4B**). Endo-GRK5 and Endo-GRK6 demonstrated both plasma membrane and endosome localization upon addition of the 2x-Fyve tag given their baseline localization to the plasma membrane.

To evaluate the signaling consequences of these GRK mutants, we utilized two previously validated CRISPR/Cas9 edited HEK293 cell lines, one devoid of GRK2 and GRK3 (ΔGRK5 and GRK6 (ΔGRK5/6)^66, 67^. We utilized these cell lines, rather than siRNA or pharmacologic strategies, to ensure complete elimination of enzymatic kinase activity. We performed immunoblotting on WT, ΔGRK5/6 HEK293 cells to determine the amount of each GRK to transfect based on WT expression levels (**Supplemental Figure 6**).

We used bioluminescence resonance energy transfer (BRET) to monitor ligand-induced β-arrestin-2-mKO recruitment to CXCR3-Renilla Luciferase II (RLucII) in either ΔGRK2/3 or ΔGRK5/6 cells transfected with the corresponding GRKs (**Figure 4C**). For CXCL10, CXCL11, and VUF11418 treatment, we surprisingly observed that rescue of WT-GRK2 decreased β-arrestin-2 recruitment while VUF10661 stimulation had no β-arrestin-2 recruitment (**Figure 4D**). Further, rescue of WT-GRK3 enhanced β-arrestin-2 recruitment for VUF11418 treatment but had no effect for other ligand conditions (**Figure 4E**). By contrast, WT-GRK5 and WT-GRK6 had little detectable effect on β-arrestin-2 recruitment for all ligands (**Supplemental Figure 7A and 7B**). These findings suggest that the WT-GRKs have isoform-specific effects on CXCR3 engagement with β-arrestin-2. These effects are conserved across ligands for some GRKs but agonist-dependent for others, indicative of the ability of different ligands to confer unique properties to GRK isoforms.

Furthermore, the effects of the mutant GRKs from the WT-GRK were ligand-specific. For example, with CXCL11, both Mem-GRK2 and Endo-GRK2 partially reduced the WT effect on β-arrestin-2 recruitment (**Figure 4D**). However, upon VUF11418 stimulation, Mem-GRK2, but not Endo-GRK2, altered the WT phenotype. A different profile was observed for GRK3. Although β-arrestin-2 recruitment was increased following rescue of WT-GRK3 with VUF11418 treatment, this effect was reduced when GRK3 was localized to the plasma membrane and completely lost when bound to endosomes (**Figure 4E**). Surprisingly, although WT-GRK3 had no detectable effect on β-arrestin-2 recruitment for CXCL10, CXCL11, and VUF10661, Mem-GRK3 decreased β-arrestin-2 recruitment for all three ligands. Whereas β-arrestin-2 recruitment was susceptible to alterations in GRK2 and GRK3 cellular localization, GRK5 and GRK6 demonstrated little dependence on cellular location for β-arrestin-2 recruitment (**Supplemental Figure 7A-7B**). These data suggest a role of location-dependent activities of the GRKs on β-arrestin-2 recruitment, where the relative direction and magnitude of change is GRK-, ligand-, and location-dependent.

Given the role of β-arrestins in promoting receptor internalization, we next assessed if individual GRKs have distinct effects on CXCR3 trafficking to endosomes. We rescued the appropriate GRKs in ΔGRK5/6 cells transfected with CXCR3-RLuc2 and 2x-Fyve-mVenus and used BRET to assess proximity to early endosomes following ligand stimulation (**Figure 4F**). While WT-GRK2 significantly decreased β engagement for CXCL10, CXCL11 and VUF11418, rescue of WT-GRK2 unexpectedly demonstrated minimal changes to CXCR3 internalization across all ligands (**Supplemental Figure 7C**). Likewise, while we observed a robust increase in β-arrestin-2 recruitment by WT-GRK3 upon VUF11418 stimulation, there was no detectable effect observed on CXCR3 internalization (**Supplemental Figure 7D**). By contrast, while WT-GRK6 showed little activity in affecting β-arrestin-2, rescue of WT-GRK6 enhanced CXCR3 internalization across ligands, except for CXCL9 (**Figure 4G**). These findings are indicative of isoform-specific roles of the GRKs in driving different GPCR signaling events. Whereas our data suggest a prominent function of GRK2 and GRK3 in regulating β-arrestin-2 recruitment to CXCR3, we found a significant role of GRK6 in mediating receptor internalization.

## GRK localization modulates isoform-specific effects on cytosolic and nuclear ERK1/2 activity in a ligand-dependent fashion

We next investigated the activation of the kinase ERK1/2 as a marker of mitogen-activated protein kinase (MAPK) signaling, a prototypical GPCR signaling pathway that can be mediated by both G proteins and β-arrestins^68, 69^. Using the ΔGRK2/3 or ΔGRK5/6 cell lines, we transfected in a previously developed BRET-based biosensor that detects ERK kinase activity in different cellular locations, specifically the nucleus or cytoplasm^42, 70^.

WT-GRK2 had little effect on cytosolic ERK activity, with minor decreases in ERK activity with CXCL9 and VUF10661 (**Figure 5A**). However, WT-GRK2 significantly inhibited nuclear ERK activity across all ligands (**Figure 5B**). In contrast, WT-GRK3 was inhibitory to both cytosolic and nuclear ERK activation for most ligands tested (**Figure 5C-5D**). WT-GRK5 had an entirely different profile, in which it diminished cytosolic ERK activity upon CXCL10 and VUF10661 stimulation but had no significant effect for these ligands on nuclear ERK activity (**Figure 5E-5F**). CXCL11 also reduced WT-GRK5-mediated cytosolic ERK phosphorylation but enhanced its activation of ERK in the nucleus. These findings support distinct roles of the GRK subtypes in regulating the activation of both cytosolic and nuclear pools of ERK. Consistent with our previous assays examining upstream signaling events, activities of GRK subtypes are specific for each ligand.

**Figure 5:**
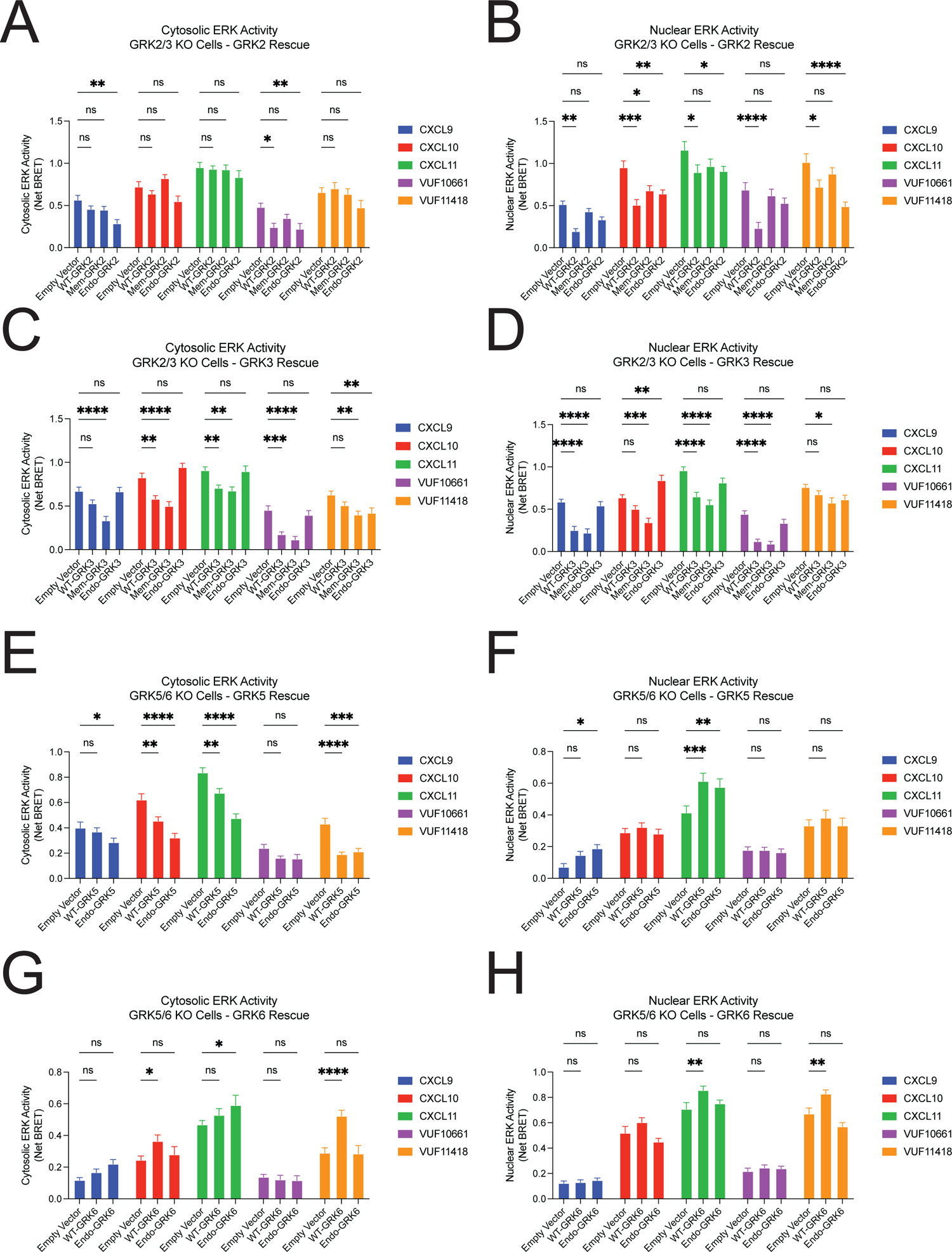
ERK activity in distinct cellular compartments in cells devoid of GRKs. Nuclear and cytoplasmic ERK activity in a ΔGRK2/3 knockout cell line upon addback of **(A and B)** GRK2 and **(C and D)** GRK3 constructs or empty vector. Nuclear and cytoplasmic ERK activity in a ΔGRK5/6 knockout cell line upon addback of **(E and F)** GRK5 and **(G and H)** GRK6 constructs or empty vector. 100nM CXCL9, CXCL10, or CXCL11, and 10µM VUF10661 or VUF11418 were used in all experiments. Data shown are the mean AUC ± SEM of n=5., *P<0.05 by two-way ANOVA with Tukey post hoc testing conducted between empty vector and other transfections conditions within each ligand.

When using the location-specific GRK mutants, Endo-GRK2 inhibited nuclear ERK activity similar to that of WT-GRK2 except at CXCL9 and VUF10661 (**Figure 5B**). For most ligands, WT-GRK3 and Mem-GRK3 significantly reduced both nuclear and cytosolic ERK activation, while Endo-GRK3 had little effect on ERK activity as compared to empty vector (**Figure 5C-5D**). Further, while WT- and Endo-GRK5 decreased cytosolic ERK activation following treatment with CXCL10, CXCL11, and VUF11418, we observed a substantial increase in nuclear ERK with CXCL11 treatment (**Figure 5E-5F**). Endo-GRK6 demonstrated little effect in modulating nuclear and cytosolic ERK activity as compared to empty vector, whereas WT-GRK6 increased ERK activation in both the nucleus and cytosol for some ligands (**Figure 5G-5H**). These data demonstrate that altering GRK localization has differential effects on the activation of nuclear and cytoplasmic ERK. By modulating the ligand, GRK, and subcellular location of the GRK, it is possible to generate numerous and highly specific patterns of ERK activity. Together, these results highlight the complex interaction between GRKs and a single ligand which can generate a diverse array of signaling responses through a GPCR.

### GRK recruitment to endosomes is observed at other GPCRs

Given our finding that biased CXCR3 agonists could promote differential GRK recruitment to the plasma membrane and endosomes, we wondered if this phenomenon is observed at other GPCRs. We selected an array of GPCRs involved in regulating a wide range of physiological processes including the β_2_-adrenergic receptor (β_2_AR), µ-opioid receptor (MOR), angiotensin II type I receptor (AT_1_R), V_2_ vasopressin receptor (V_2_R) and atypical chemokine receptor 3 (ACKR3). We then probed GRK2, 3, 5, and 6 recruitment to the plasma membrane and endosomes following ligand stimulation using the aforementioned Nano-BiT complementation assay. Cells were transfected with WT receptor, the indicated GRK-LgBiT and either CD8⍰-SmBiT for the plasma membrane or 2xFyve-SmBiT for the endosome (**Figure 6**).

**Figure 6:**
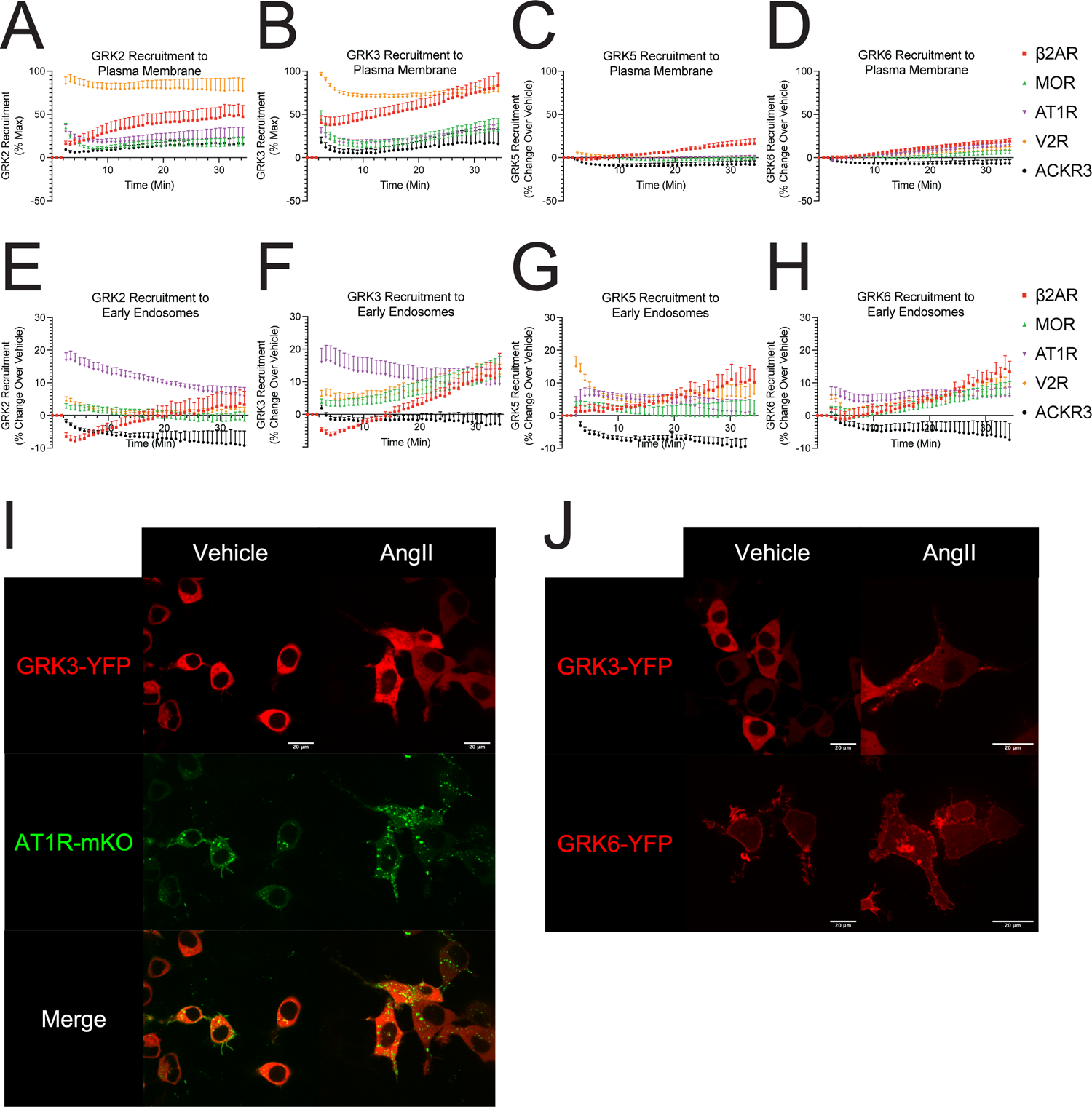
GRK recruitment to plasma membrane and endosomes at other GPCRs. Recruitment of **(A)** GRK2, **(B)** GRK3, **(C)** GRK5, and **(D)** GRK6 to the plasma membrane upon stimulation of each specified receptor with a single dose of the indicated ligand in HEK293 cells. Recruitment of **(E)** GRK2, **(F)** GRK3, **(G)** GRK5, and **(H)** GRK6 to the endosome upon stimulation of each specified receptor with a single dose of the indicated ligand in HEK293 cells. **(I)** Confocal microscopy of mKO tagged angiotensin II type I receptor (AT1R-mKO) and GRK3-YFP 45 minutes after addition of vehicle or angiotensin II (AngII). **(J and K)** Confocal microscopy of untagged angiotensin II type I receptor (AT1R-mKO) and GRK3-YFP or GRK6-YFP 30 minutes after addition of vehicle or angiotensin II (AngII). The β_2_-adrenergic receptor (B2AR) was stimulated with 10µM isoproterenol, the µ-opioid receptor (MOR) was stimulated with 10µM DAMGO, the angiotensin II type I receptor (AT1R) was stimulated with 10µM angiotensin II, the V_2_ vasopressin receptor (V2R) was stimulated with 1µM arginine vasopressin, and the atypical chemokine receptor 3 (ACKR3) was stimulated with 10µM of WW36. Data shown are the mean ± SEM of n=5 and are normalized luminescence change over vehicle. Confocal microscopy images are representative of n=3.

We demonstrate that the receptors studied differ in their ability to induce GRK recruitment to the plasma membrane. For example, stimulation of the V_2_R produced the most robust recruitment of GRK2 and GRK3 to the plasma membrane, whereas stimulation of the β_2_AR induced the greatest recruitment of GRK5 and GRK6 to the plasma membrane (**Figure 6A-6D**). While GRK5 and GRK6 are constitutively membrane localized, this may represent a change in the membrane distribution of these GRKs as CD8⍰ can be found in specific membrane microdomains^71, 72^.

Consistent with our data at CXCR3, we observed significant recruitment of the GRKs to endosomes for many of the receptors tested, with distinct patterns when compared to plasma membrane recruitment (**Figure 6E-6H**). GRK2 and GRK3 were significantly recruited to endosomes upon activation of the AT_1_R (**Figure 6E and 6F**). The β_2_AR, MOR, and V_2_R all demonstrated the ability to recruit GRK2 and or GRK3 to endosomes, albeit to different magnitudes and with different kinetic tracings. GRK5 and GRK6 similarly demonstrated recruitment to endosomes across many receptors in a pattern that was different than that observed at the plasma membrane. (**Figure 6C-6D and 6G-6H)**. We further validated these findings at the AT_1_R using confocal microscopy. Following treatment with angiotensin II (AngII), we observed colocalization of GRK3-YFP and AT1R-mKO in endosomes (**Figure 6I**). We repeated these experiments using an untagged AT1R and observed formation of both GRK3-YFP and GRK6-YFP containing endosomes following treatment with AngII (**Figure 6J**). These data suggest that trafficking of GRKs to endosomes is a conserved signaling mechanism for many GPCRs, and that distinct patterns of GRK engagement with GPCRs at the plasma membrane and endosomes may facilitate signaling specificity and promote biased responses.

## DISCUSSION

While almost all GPCRs are known to interact with G proteins, GRKs, and β-arrestins, these interactions have primarily been studied at the plasma membrane. While recent data demonstrates that G proteins and β-arrestins can signal from endosomes^73–75^, it was unknown if GRKs exhibited similar behavior. Here, we provide, to our knowledge, some of the first evidence that GRKs can traffic to endosomes following agonist stimulation of a GPCR. Additionally, using both endogenous and synthetic CXCR3 agonists, we show that the pattern of GRK recruitment to the plasma membrane differs significantly from that observed at endosomes. Whereas CXCL11 promoted robust GRK2 recruitment to CXCR3 at the plasma membrane, this ligand did not produce any detectable engagement of GRK2 in endosomes. Similarly, stimulation of the V_2_R promotes significantly more GRK2 and GRK3 recruitment to the plasma membrane than the AT_1_R, but this pattern is reversed in endosomes. It is possible that subcellular localization of the GRKs may contribute to the ability of these GPCRs to initiate spatially distinct signaling patterns. Elucidation of the unique roles of individual GRK subtypes on overall GPCR signaling has received appreciable attention recently, in large measure due to the potential contribution of single GRKs to a wide range of pathophysiological processes. The dysregulation of GRK2 has been linked to the progression of mitochondrial lesions in Alzheimer’s disease, cardiovascular dysfunction, pathological angiogenesis, and chronic inflammation, among others^76–80^.

Using biased ligands, we observed that G protein activation is a critical mechanism underlying the biased recruitment of GRKs to the plasma membrane at CXCR3. These findings are consistent with previous reports which demonstrate that GRK2 and GRK3 are recruited to GPCRs via their PH domain recognizing free Gβγ following heterotrimeric G protein activation^29, 57, 81^. However, using two different CRISPR/Cas9 GRK KO cell lines, we demonstrate that GRK2 and GRK3 can be recruited to CXCR3 in independent of G protein activation. Interestingly, the relative amount of G protein-independent GRK recruitment observed was dependent on the ligand, GRK, and cell type used. It is possible that these observations are a consequence of using genetically modified HEK293 cells. However, other work has demonstrated a similar phenomenon at the dopamine D2 receptor, which can recruit GRK2 in the absence of G protein activation^82^. We build upon this previous work as our data suggest that a single GPCR may use both G protein-dependent and independent mechanisms to recruit GRKs to the plasma membrane, depending on the ligand used to activate the receptor. We finally explored the contributions of GRK subcellular localization to biased GPCR signaling by developing location-based GRK mutants and characterizing CXCR3 signaling using GRK KO cell lines. We demonstrate that modifications to GRK cellular localization can alter the receptor’s ability to recruit β-arrestin 2, internalize, and activate ERK, depending on the specific ligand and GRK being studied. Thus, GRK subtype functionality appears to be not only driven by distinctions between the isoforms themselves, but also by unique properties conferred to each GRK by different ligands.

Receptor internalization and recycling was initially considered a mechanism of signal attenuation and re-sensitization in response to ligand stimulation^83, 84^. Specifically, β-arrestins sterically inhibit GPCRs to prevent further G protein activation while simultaneously promoting receptor internalization to prevent further ligand binding. However, recent studies demonstrate that many GPCRs can continue to signal through both G proteins and β-arrestins from endosomes. For example, previous work at the β_2_AR showed that endosomal cAMP production is required to generate a complete transcriptional response^41^. Subsequent work has showed that signaling from endosomes also impacts the global phosphoproteome^40^. Many studies support the paradigm of *location bias*, and it is now appreciated that endocytosis does not solely serve as a mechanism to abrogate GPCR signaling; rather, it also can be a critical component to achieving maximal GPCR signaling^38, 85–88^. Additionally, recent work demonstrates the therapeutic potential of pharmacologically targeting GPCR signaling from endosomes at the neurokinin 1 receptor, MOR, and other receptors^89–92^.

Importantly, recent studies have observed that a specific ligand:GPCR complex can generate signaling pathways from endosomes that are distinct from those generated at the plasma membrane^39, 41, 88^. It is likely that signaling from endosomes, and other subcellular locations, is one mechanism by which a GPCR can generate diverse signaling outputs in response to biased ligands. We recently demonstrated that the biased agonists of CXCR3 demonstrate different relative amounts of G protein and β-arrestin signaling at the plasma membrane and endosome, providing evidence that the degree of biased agonism observed at a GPCR depends on where in the cell the biased signaling is measured^42^. However, the mechanisms underlying this location-specific signaling are incompletely understood.

Notably, changes in a given readout of CXCR3 signaling were not necessarily predictive of the alterations for other signaling events canonically viewed as sequential. For example, although differential GRK6 localization had little detectable effect on the ability of CXCR3 to engage β-arrestin 2, GRK6 increased CXCR3 internalization across all ligands. These results are consistent with GRKs demonstrating function beyond receptor phosphorylation and increasing affinity of β-arrestin for a GPCR. Recent research demonstrates that the GRKs can modulate β-arrestin conformation which is tightly associated with its ability to scaffold other signaling effector^67, 93–95^. Additionally, GRKs are known to phosphorylate other non-GPCR receptors such as receptor tyrosine kinases and toll-like receptors, transcription factors, and more^28^. GRKs are also known to have kinase-independent functionality. For example, previous research demonstrated that down-regulation of GRK2 leads to significant impairment in zebrafish development. However, normal development can be rescued by expressing a kinase-dead mutant of GRK2. The authors discovered that GRK2 forms an interaction with cyclin B1 regulator patched homolog 1 (PTCH1), independent of its kinase activity, which was required necessary for normal embryogenesis^96^. Similar work has demonstrated that GRK5 can inhibiting NFκB signaling through nuclear accumulation of IκB^97^.

Our findings highlight the importance of the GRKs in initiating biased responses at GPCRs and demonstrate the complex signaling that can be achieved through a single GPCR. These data support the phosphorylation barcode hypothesis by demonstrating that differential GPCR engagement with the GRKs can generate multiple, agonist-specific, signaling pathways. Importantly, we observed that this complex signaling exists at different compartments within the cell. Together, our data demonstrate the therapeutic promise and simultaneous complexity of drugging GPCRs and GPCR effectors given the vast diversity of signaling that can be achieved within this receptor family. The molecular determinants which promote GRK translocation to endosomes are still unclear at this time, and the complex interconnection between ligand and GRK subtype bias has not been fully resolved. It will be important to determine the mechanisms by which individual GRKs modulate signaling at different locations. Deconvoluting the structural elements that exist between a GPCR and GRK are critical to understanding the biochemical basis for biased signaling observed across different cellular compartments.

## ACKNOWLEDGEMENTS

We thank Asuka Inoue (Tohoku University, Japan), Robert J. Lefkowitz (Duke University, USA), Christopher Cole Honeycutt (National Institutes of Health, USA), and Noelia Boldizsar (National Institutes of Health, USA) for reagents, guidance, and thoughtful feedback throughout this work; N. Nazo for laboratory assistance. Funding: This work was supported by T32GM007171 (D.S.E.), the Duke Medical Scientist Training Program (D.S.E.), AHA 20PRE35120592 (D.S.E.), 1R01GM122798 (S.R.), Burroughs Wellcome Career Award for Medical Scientists (S.R.). Graphical figures were created with BioRender.

## AUTHOR CONTRIBUTIONS

Conceptualization, J.G., D.S.E., S.R.; Methodology, J.G., D.S.E., C.H., S.R.; Investigation, J.G., D.S.E., C.H., I.C.; Writing — Original Draft, J.G., D.S.E., C.H.; Writing – Reviewing & Editing, J.G., D.S.E., C.H., N.B., C.C.H., U.P, and S.R.; Visualization, J.G., D.S.E., C.H, and S.R.; Supervision and Funding Acquisition, S.R.

## METHODS

### RESOURCE AVAILABILITY

#### Lead Contact

Further information and requests for resources and reagents should be directed to and will be fulfilled by the lead contact, Sudarshan Rajagopal.

#### Materials Availability

All plasmids generated in this study will be distributed upon request.

#### Data and Code Availability

All data reported in this paper will be shared by the lead contact upon request.

## EXPERIMENTAL MODEL AND SUBJECT DETAILS

### Bacterial Strains

XL-10 Gold ultracompetent E. coli (Agilent) were used to express all constructs used in this manuscript.

### Cell Lines

Human Embryonic Kidney (HEK293) cells were grown in minimum essential media (MEM) supplemented with 10% fetal bovine serum (FBS) and 1% penicillin/streptomycin at 37°C and 5% CO2. ΔGRK2/3, ΔGRK5/6, ΔG6, and ΔG7 CRISPR/Cas9 KO HEK293 cells were provided by Asuka Inoue, Tohoku University, Japan, and validated as previously described^47, 62, 65^.

## METHOD DETAILS

### Generation of Constructs

Construct cloning was performed using conventional techniques such as restriction enzyme/ligation methods. Linkers between the fluorescent proteins or luciferases and the cDNAs for receptors, transducers, or other proteins were flexible and ranged between 2 and 18 amino acids. Fluorescence resonance energy transfer (FRET) based ERK1/2 biosensors previously published (Harvey et al., 2008) were used to generate BRET versions of these sensors by removing the N-terminal mCerulean through restriction digest and inserting a nanoluciferase. We utilized the 2x-Fyve targeting sequence (RKHHCRACG) from the hepatocyte growth factor-regulated tyrosine kinase substrate to target labeled constructs to endosomes. We utilized the membrane targeting sequence (MGCIKSKGKDS) from Lyn kinase to target labeled constructs to the plasma membrane. Location tags were attached to the N-terminus of all constructs, followed by a flexible amino acid linker.

### Cell Culture and Transfection

For luminescence-based assays, HEK293 cells were transiently transfected with an optimized calcium phosphate protocol as previously described unless otherwise indicated^82^. In the calcium phosphate transfection method, cell culture media was replaced 30 minutes prior to transfection. Plasmid constructs were suspended in water to a final volume of 90µL. 10μL of 2.5 M calcium chloride was added to the plasmid constructs and mixed. 100μL of 2x HEPES-buffered saline solution (10mM D-Glucose, 40mM HEPES, 10 mM potassium chloride, 270 mM sodium chloride, 1.5 mM disodium hydrogen phosphate dihydrate) was added to the solution, allowed to incubate for two minutes, and subsequently added to the cells. For BRET based assays, luminescence-based assays in G protein KO cell lines, and confocal microscopy, cells were transiently transfected using polyethylenimine (PEI). In the PEI transfection method, cell culture media was replaced 30 minutes prior to transfection. Plasmid constructs were suspended in Opti-MEM (Gibco) to a final volume of 100 μL and, in a separate tube, PEI at a concentration of 1 mg/mL was added to Opti-MEM to a final volume of 100μL. For experiments in this manuscript, 3μL of PEI was used per 1μg of plasmid DNA. After 5 minutes, the 100μL PEI solution was added to the 100μL DNA solution, gently mixed, and allowed to incubate at room temperature for 10-15 minutes, after which the mixture was added to the cells.

### BRET and Split Luciferase Assays

For all BRET and Split Luciferase assays, HEK293 cells seeded in 6 well plates (∼750,000 cells/well) were transiently transfected with the appropriate constructs using the calcium phosphate or PEI method previously described. GRK recruitment was assessed using a NanoBiT complementation assay, where GRK2, GRK3, GRK5, or GRK6 were tagged with a C-terminal LgBiT and transfected with either CXCR3-smBiT (receptor), wild-type CXCR3 and 2x-Fyve-SmBiT (endosome), or wild-type CXCR3 and CD8⍰-SmBiT (plasma membrane). The role of G protein activation in GRK2/3 recruitment was assessed using PEI transfection of CXCR3-SmBiT and the indicated GRK-LgBiT, with rescue of Gi1 in ΔG7 GKO cell lines and pertussis toxin (PTX) treatment at a final concentration of 200ng/mL in ΔG6 GKO cells.

To examine role of GRK subcellular localization on effector engagement and downstream signaling, we assessed β-arrestin 2 recruitment and CXCR3 internalization using the wild-type GRKs, membrane-bound GRK2 or 3 constructs, or endosome-bound GRK2, 3, 5, or 6. Rescue of GRK2 or 3 was performed in GRK2/3 KO cells while rescue of GRK5 or 6 was performed in GRK 5/6 KO cells. For β-arrestin-2 recruitment assays, cells were transfected with the indicated WT- or mutant GRK, CXCR3-RLuc2, and β-arrestin-2-mKO. CXCR3 internalization assays used the indicated WT- or mutant GRK with CXCR3-RLuc2 and a 2x-Fyve-mVenus to assess proximity to the early endosome.

Location-specific BRET-biosensors of ERK activity were transfected using PEI. This ERK biosensor consists of a target ERK substrate that, following phosphorylation by activated phosphorylated ERK, binds to a phosphorylation binding domain, causing a conformational change in the biosensor and subsequent change in BRET efficiency^42^.

Twenty-four hours after transfection, cells were washed with phosphate buffered saline (PBS), collected with trypsin, and plated onto clear-bottomed, white-walled, Costar 96-well plates at 50,000 to 100,000 cells/well in BRET medium (clear minimum essential medium (Gibco) supplemented with 2% FBS, 10 mM HEPES, 1x GlutaMax (Gibco), and 1x Antibiotic-Antimycotic (Gibco)).

The following day, media was removed, and cells were incubated at 37°C with 80 μL of HBSS supplemented with 20mM HEPES, and 3µM coelenterazine h for all BRET or NanoBiT complementation assays (Cayman Chemical, Ann Arbor, MI and Nanolight Technology, Pinetop, AZ) for 10 to 15 minutes. For all BRET assays, a standard 480nm RLuc emission filter and 530nm (for experiment using mVenus) or custom 542-nm (for experiments using mKO) long pass filter was utilized (Chroma Technology Co., Bellows Falls, VT). Cells were stimulated with either vehicle control (HBSS with 20 mM HEPES) or the indicated concentration of ligand. For NanoBiT complementation and BRET experiments, three initial reads were taken prior to the addition of ligand to quantify baseline luminescence or BRET signal before adding ligand. The change in luminescence after ligand stimulation was subsequently normalized to vehicle treatment. For BRET experiments, the BRET ratio was calculated by dividing the acceptor signal by the luminescent signal, and a net BRET ratio was calculated by normalizing to vehicle treatment. Plates were read with a BioTek Synergy Neo2 plate reader or Berthold Mithras LB 940 set at 37°C. All readings were performed using a kinetic protocol.

### Confocal Microscopy

HEK293 cells were plated on 35 mm glass bottomed dishes (MatTek Corporation, Ashland, MA) and transiently transfected using PEI with the listed constructs. To evaluate the baseline expression of wild-type GRK2-YFP, GRK3-YFP, GRK5-YFP, and GRK6-YFP, or the Lyn-tagged or 2xFyve-tagged indicated GRK-YFP, cells were imaged forty-eight hours after transfection. To assess ligand-induced GRK localization, forty-eight hours after transfection, the cells were washed once with PBS and then serum starved for one hour. The cells were subsequently treated with a control of serum free media or the listed chemokine at 100nM or listed VUF compound at 10 μM for forty-five minutes at 37°C. Cells were then imaged with a Zeiss CSU-X1 spinning disk confocal microscope using the corresponding lasers for YFP (480nm excitation) or mCerulean (433nm excitation). Images were analyzed using ImageJ (NIH, Bethesda, MD) and false colors were applied for clarity.

### Immunoblotting

Immunoblotting was performed as described previously^43^. ΔGRK2/3 and ΔGRK5/6 cells seeded in 6 well plates were transiently transfected with increasing amounts of GRK using the calcium phosphate transfection method. 24 hours after transfection, cells were serum starved in minimum essential medium supplemented with 0.01% bovine serum albumin (BSA) and 1% penicillin/streptomycin for 16 hours. The cells were then washed with ice cold PBS, and lysed in ice cold RIPA buffer supplemented with protease inhibitors (Phos-STOP (Roche), cOmplete EDTA free (Sigma)). The samples were rotated at 4°C for forty-five minutes and cleared of insoluble debris by centrifugation at 17,000g at 4°C for 15 minutes, after which the supernatant was collected. Protein was resolved on SDS-10% polyacrylamide gels, transferred to nitrocellulose membranes, and immunoblotted with the indicated primary antibody overnight at 4°C. GRK2 (Santa Cruz, #sc-13143), GRK3 (Cell Signaling Technology, #80362), GRK5 (Santa Cruz, #sc-518005), and GRK6 (Cell Signaling Technology, #5878) antibodies were used to compare rescued GRK expression level in Δ GRK5/6 cells to GRK expression levels WT HEK293 cells. Immunoblots were normalized using an alpha-tubulin antibody (Sigma, #T5168). Horseradish peroxidase-conjugated anti-rabbit-IgG or anti-mouse-IgG were used as secondary antibodies. The nitrocellulose membranes were imaged by SuperSignal enhanced chemiluminescent substrate (Thermo Fisher) using a ChemiDoc MP Imaging System (Bio-Rad).

### CXCR3 Ligands

Recombinant Human CXCL9, CXCL10, and CXCL11 (PeproTech) were diluted according to the manufacturer’s specifications, and aliquots were stored at −80°C until needed for use. VUF10661 (Sigma-Aldrich) and VUF11418 (Aobius) were reconstituted in dimethyl sulfoxide (DMSO) and stored at −20°C in a desiccator cabinet.

## QUANTIFICATION AND STATISTICAL ANALYSIS

Data were analyzed in Excel (Microsoft, Redmond, WA) and graphed in Prism 9.0 (GraphPad, San Diego, CA). Dose-response curves were fitted to a log agonist versus stimulus with three parameters (span, baseline, and EC50), with the minimum baseline corrected to zero using Prism 9.0. Statistical tests were performed using a one or two-way ANOVA followed by Tukey’s multiple comparison’s test when comparing treatment conditions. When comparing ligands or treatment conditions in concentration-response assays or time-response assays, a two-way ANOVA of ligand and concentration or ligand and AUC, respectively, was conducted. For experiments using mutant GRKs, a two-way ANOVA was performed followed by Tukey’s multiple comparison’s test when comparing transfection conditions within a ligand. If a significant interaction effect was observed (*P* < 0.05), then comparative two-way ANOVAs between individual experimental conditions were performed. Further details of statistical analysis and replicates are included in the figure legends. Lines represent the mean, and error bars signify the SEM, unless otherwise noted. Further details of statistical analysis and replicates are included in the figure captions. Experiments were not randomized, and investigators were not blinded to treatment conditions. Critical plate-based experiments were independently replicated by at least two different investigators when feasible.

## KEY RESOURCES TABLE

| REAGENT OR RESOURCE | SOURCE | IDENTIFIER |
| --- | --- | --- |
| **Antibodies** |  |  |
| Donkey polyclonal anti-rabbit IgG peroxidase conjugated | Rockland | Cat#611-7302 |
| Sheep polyclonal anti-mouse IgG peroxidase conjugated | Rockland | Cat#610-603-002 |
| Mouse monoclonal anti-GRK2 IgG | Santa Cruz | Cat# sc-13143 |
| Rabbit monoclonal anti-GRK3 IgG | Cell Signaling Technology | Cat#80362 |
| Mouse monoclonal anti-GRK5 IgG | Santa Cruz | Cat# sc-518005 |
| Rabbit monoclonal anti-GRK6 IgG | Cell Signaling Technology | Cat#5878 |
| **Bacterial Strains** |  |  |
| XL 10-Gold Ultracompetent E. Coli | Agilent | Cat#200315 |
| **Chemicals, peptides, and recombinant proteins** |  |  |
| Recombinant Human CXCL9 | Peprtech | Cat#300-26 |
| Recombinant Human CXCL10 | Peprtech | Cat#300-12 |
| Recombinant Human CXCL11 | Peprtech | Cat#300-46 |
| VUF10661 | Sigma-Aldrich | Cat#SML0803 |
| VUF11418 | Aobius | Cat#AOB<sub>3998</sub> |
| Angiotensin II human | Sigma-Aldrich | Cat#A9525 |
| Isoproterenol hydrochloride | Sigma-Aldrich | Cat#I6504 |
| [Arg<sup>8</sup>]-Vasopressin | Sigma-Aldrich | Cat#V9879 |
| DAMGO | MedChem Express | Cat#HY-P0210 |
| WW36 | United Therapeutics |  |
| GlutaMax | Gibco | Cat#35050061 |
| Antibiotic-Antimycotic | Gibco | Cat#15240062 |
| Coelenterazine h | Cayman Chemical | Cat#16894 |
| Coelenterazine h | NanoLight Technology | Cat#301 |
| **Cell Lines** |  |  |
| Human: 293 | ATCC | Cat#CRL-1573 |
| Human: 293T | ATCC | Cat#CRL-3216 |
| Human: 293T GRK2/3 Knock Out | Asuka Inoue | <sup>47</sup> |
| Human: 293T GRK5/6 Knock Out | Asuka Inoue | <sup>47</sup> |
| Human: 293T ΔG6 G protein Knock Out | Asuka Inoue | <sup>62</sup> |
| Human: 293T ΔG7 G protein Knock Out | Asuka Inoue | <sup>65</sup> |

### SUPPLEMENTAL FIGURE TITLES AND LEGENDS

**Supplemental Figure 1: Confocal images of GRK2, GRK3, GRK5, and GRK6**. Confocal microscopy images of HEK293 cells transfected with CXCR3-mCerulean (shown in green) and YFP-tagged GRK2, GRK3, GRK5, or GRK6 (shown in red). False colors are applied for clarity. Images were taken in the absence of ligand stimulation. Confocal microscopy images are representative of n=3.

**Supplemental Figure 2: Confocal microscopy of GRK2 and GRK3 following activation of CXCR3.** Confocal microscopy images of HEK293 cells transfected with CXCR3-mCerulean (shown in green) and **(A)** GRK2-YFP and **(B)** GRK3-YFP (shown in red). False colors are applied for clarity. Images were taken 45 minutes after stimulation with listed treatment. Confocal microscopy images are representative of n=3.

**Supplemental Figure 3: Confocal microscopy of GRK5 and GRK6 following activation of CXCR3.** Confocal microscopy images of HEK293 cells transfected with CXCR3-mCerulean (shown in green) and **(A)** GRK5-YFP and **(B)** GRK6-YFP (shown in red). False colors are applied for clarity. Images were taken 45 minutes after stimulation with listed treatment. Confocal microscopy images are representative of n=3.

**Supplemental Figure 4:** Kinetic data of GRK2 and GRK3 recruitment in Δ**G6 cells.** Kinetic data of **(A-E)** GRK2 and **(F-J)** GRK3 recruitment to CXCR3 with and without 200ng/mL pertussis toxin treatment (PTX) in ΔG6 cells treated with 100nM CXCL9, CXCL10, or CXCL11, and 10µM VUF10661 or VUF11418. GRK recruitment was measured using a NanoBiT complementation assay involving LgBiT-tagged GRK2 or GRK3, and CXCR3-SmBiT. Data shown are the mean ± SEM of n=5. *P<0.05 by two-way ANOVA with Tukey post hoc testing between treatment conditions at a single GRK:ligand combination. Dose response curves are normalized to maximum signal observed across all ligands.

**Supplemental Figure 5: Dose response and kinetic data of GRK2 and GRK3 recruitment in** Δ**G7 cells.** Dose response data of **(A-E)** GRK2 and **(K-O)** GRK3 recruitment to CXCR3 and kinetic data of **(F-J)** GRK2 and **(P-T)** GRK3 recruitment to CXCR3 in ΔG7 cells transfected with empty vector (pcDNA 3.1) or G⍰i, treated with 100nM CXCL9, CXCL10, or CXCL11, and 10µM VUF10661 or VUF11418. GRK recruitment was measured using a NanoBiT complementation assay involving LgBiT-tagged GRK2 or GRK3, and CXCR3-SmBiT. Data shown are the mean ± SEM of n=5. *P<0.05 by two-way ANOVA with Tukey post hoc testing between transfections conditions at a single GRK:ligand combination. Dose response curves are normalized to maximum signal observed across all ligands.

**Supplemental Figure 6: Immunoblots of rescued GRK expression in** Δ**GRK5/6 cells.** Immunoblots demonstrating increasing amounts of transfected **(A)** GRK2, **(B)** GRK3, **(C)** GRK5, and **(D)** GRK6 in corresponding parental KO cell lines as compared to wild-type HEK293 cells.

**Supplemental Figure 7:** β**-arrestin recruitment and CXCR3 internalization data in** Δ**GRK2/3 and** Δ**GRK5/6 cells.** β-arrestin 2 recruitment to CXCR3 as measured by BRET in a ΔGRK5/6 cells following addition of addback of **(A)** GRK5 and **(B)** GRK6 constructs or empty vector. CXCR3 internalization as measured by BRET in ΔGRK2/3 cells upon addback of **(C)** GRK2 and **(D)** GRK3 constructs or empty vector, ΔGRK5/6 cells upon addback of **(E)** GRK5. 100nM CXCL9, CXCL10, or CXCL11, and 10µM VUF10661 or VUF11418 were used in all experiments. *P<0.05 by two-way ANOVA with Tukey post hoc testing conducted between empty vector and other transfections conditions within each ligand.

## REFERENCES

1. Bockaert, J. & Pin, J.P. Molecular tinkering of G protein-coupled receptors: an evolutionary success. EMBO J 18, 1723–9 (1999).

2. Santos, R. et al. A comprehensive map of molecular drug targets. Nat Rev Drug Discov 16, 19–34 (2017).

3. Zhang, Y., Devries, M.E. & Skolnick, J. Structure modeling of all identified G protein-coupled receptors in the human genome. PLoS Comput Biol 2, e13 (2006).

4. Eiger, D.S., Pham, U., Gardner, J., Hicks, C. & Rajagopal, S. GPCR systems pharmacology: a different perspective on the development of biased therapeutics. Am J Physiol Cell Physiol 322, C887–C895 (2022).

5. Reiter, E. & Lefkowitz, R.J. GRKs and beta-arrestins: roles in receptor silencing, trafficking and signaling. Trends Endocrinol Metab 17, 159–65 (2006).

6. Eishingdrelo, H. & Kongsamut, S. Minireview: Targeting GPCR Activated ERK Pathways for Drug Discovery. Curr Chem Genom Transl Med 7, 9–15 (2013).

7. Tian, X., Kang, D.S. & Benovic, J.L. β-arrestins and G protein-coupled receptor trafficking. Handb Exp Pharmacol 219, 173–86 (2014).

8. Caron, M.G. & Barak, L.S. A Brief History of the beta-Arrestins. Methods Mol Biol 1957, 3–8 (2019).

9. Rajagopal, S. et al. Quantifying ligand bias at seven-transmembrane receptors. Mol Pharmacol 80, 367–77 (2011).

10. Shenoy, S.K. Arrestin interaction with E3 ubiquitin ligases and deubiquitinases: functional and therapeutic implications. Handb Exp Pharmacol 219, 187–203 (2014).

11. Kenakin, T. & Christopoulos, A. Signalling bias in new drug discovery: detection, quantification and therapeutic impact. Nat Rev Drug Discov 12, 205–16 (2013).

12. Smith, J.S., Lefkowitz, R.J. & Rajagopal, S. Biased signalling: from simple switches to allosteric microprocessors. Nat Rev Drug Discov 17, 243–260 (2018).

13. Eiger, D.S., Boldizsar, N., Honeycutt, C.C., Gardner, J. & Rajagopal, S. Biased agonism at chemokine receptors. Cell Signal 78, 109862 (2021).

14. Yang, Z. et al. Phosphorylation of G Protein-Coupled Receptors: From the Barcode Hypothesis to the Flute Model. Mol Pharmacol 92, 201–210 (2017).

15. Nobles, K.N. et al. Distinct phosphorylation sites on the beta(2)-adrenergic receptor establish a barcode that encodes differential functions of beta-arrestin. Sci Signal 4, ra51 (2011).

16. Xiao, K. et al. Global phosphorylation analysis of beta-arrestin-mediated signaling downstream of a seven transmembrane receptor (7TMR). Proc Natl Acad Sci U S A 107, 15299–304 (2010).

17. Kaya, A.I., Perry, N.A., Gurevich, V.V. & Iverson, T.M. Phosphorylation barcode-dependent signal bias of the dopamine D1 receptor. Proceedings of the National Academy of Sciences 117, 14139–14149 (2020).

18. Latorraca, N.R. et al. How GPCR Phosphorylation Patterns Orchestrate Arrestin-Mediated Signaling. Cell 183, 1813–1825 e18 (2020).

19. Dwivedi-Agnihotri, H. et al. Distinct phosphorylation sites in a prototypical GPCR differently orchestrate beta-arrestin interaction, trafficking, and signaling. Sci Adv 6(2020).

20. Liggett, S.B. Phosphorylation barcoding as a mechanism of directing GPCR signaling. Sci Signal 4, pe36 (2011).

21. Premont, R.T. et al. The GRK4 subfamily of G protein-coupled receptor kinases. Alternative splicing, gene organization, and sequence conservation. J Biol Chem 274, 29381–9 (1999).

22. Lodowski, D.T., Tesmer, V.M., Benovic, J.L. & Tesmer, J.J. The structure of G protein-coupled receptor kinase (GRK)-6 defines a second lineage of GRKs. J Biol Chem 281, 16785–93 (2006).

23. Packiriswamy, N. & Parameswaran, N. G-protein-coupled receptor kinases in inflammation and disease. Genes Immun 16, 367–77 (2015).

24. Katritch, V., Cherezov, V. & Stevens, R.C. Diversity and modularity of G protein-coupled receptor structures. Trends Pharmacol Sci 33, 17–27 (2012).

25. Komolov, K.E. & Benovic, J.L. G protein-coupled receptor kinases: Past, present and future. Cell Signal 41, 17–24 (2018).

26. Chaudhary, P.K. & Kim, S. The GRKs Reactome: Role in Cell Biology and Pathology. Int J Mol Sci 22(2021).

27. Matthees, E.S.F., Haider, R.S., Hoffmann, C. & Drube, J. Differential Regulation of GPCRs-Are GRK Expression Levels the Key? Front Cell Dev Biol 9, 687489 (2021).

28. Gurevich, E.V., Tesmer, J.J., Mushegian, A. & Gurevich, V.V. G protein-coupled receptor kinases: more than just kinases and not only for GPCRs. Pharmacol Ther 133, 40–69 (2012).

29. Touhara, K., Inglese, J., Pitcher, J.A., Shaw, G. & Lefkowitz, R.J. Binding of G protein beta gamma-subunits to pleckstrin homology domains. J Biol Chem 269, 10217–20 (1994).

30. Thiyagarajan, M.M. et al. A predicted amphipathic helix mediates plasma membrane localization of GRK5. J Biol Chem 279, 17989–95 (2004).

31. Jiang, X., Benovic, J.L. & Wedegaertner, P.B. Plasma membrane and nuclear localization of G protein coupled receptor kinase 6A. Mol Biol Cell 18, 2960–9 (2007).

32. Doll, C. et al. Deciphering micro-opioid receptor phosphorylation and dephosphorylation in HEK293 cells. Br J Pharmacol 167, 1259–70 (2012).

33. Butcher, A.J. et al. Differential G-protein-coupled receptor phosphorylation provides evidence for a signaling bar code. J Biol Chem 286, 11506–18 (2011).

34. Fredericks, Z.L., Pitcher, J.A. & Lefkowitz, R.J. Identification of the G protein-coupled receptor kinase phosphorylation sites in the human beta2-adrenergic receptor. J Biol Chem 271, 13796–803 (1996).

35. Lin, F.T., Daaka, Y. & Lefkowitz, R.J. beta-arrestins regulate mitogenic signaling and clathrin-mediated endocytosis of the insulin-like growth factor I receptor. J Biol Chem 273, 31640–3 (1998).

36. Jensen, D.D. et al. Neurokinin 1 receptor signaling in endosomes mediates sustained nociception and is a viable therapeutic target for prolonged pain relief. Sci Transl Med 9(2017).

37. Sakamoto, H. et al. Expression of G protein-coupled receptor-30, a G protein-coupled membrane estrogen receptor, in oxytocin neurons of the rat paraventricular and supraoptic nuclei. Endocrinology 148, 5842–50 (2007).

38. Irannejad, R., Tsvetanova, N.G., Lobingier, B.T. & von Zastrow, M. Effects of endocytosis on receptor-mediated signaling. Curr Opin Cell Biol 35, 137–43 (2015).

39. Tsvetanova, N.G., Irannejad, R. & von Zastrow, M. G protein-coupled receptor (GPCR) signaling via heterotrimeric G proteins from endosomes. J Biol Chem 290, 6689–96 (2015).

40. Tsvetanova, N.G. et al. Endosomal cAMP production broadly impacts the cellular phosphoproteome. J Biol Chem 297, 100907 (2021).

41. Tsvetanova, N.G. & von Zastrow, M. Spatial encoding of cyclic AMP signaling specificity by GPCR endocytosis. Nat Chem Biol 10, 1061–5 (2014).

42. Eiger, D.S., et al. Location bias contributes to functionally selective responses of biased CXCR3 agonists. *bioRxiv*, 2022.01.13.476255 (2022).

43. Smith, J.S. et al. Biased agonists of the chemokine receptor CXCR3 differentially control chemotaxis and inflammation. Sci Signal 11(2018).

44. Zheng, K. et al. Biased agonists of the chemokine receptor CXCR3 differentially signal through Galphai:beta-arrestin complexes. Sci Signal 15, eabg5203 (2022).

45. Colvin, R.A., Campanella, G.S., Manice, L.A. & Luster, A.D. CXCR3 requires tyrosine sulfation for ligand binding and a second extracellular loop arginine residue for ligand-induced chemotaxis. Mol Cell Biol 26, 5838–49 (2006).

46. Groom, J.R. & Luster, A.D. CXCR3 in T cell function. Exp Cell Res 317, 620–31 (2011).

47. Kawakami, K. et al. Heterotrimeric Gq proteins act as a switch for GRK5/6 selectivity underlying beta-arrestin transducer bias. Nat Commun 13, 487 (2022).

48. Scholten, D.J. et al. Pharmacological characterization of a small-molecule agonist for the chemokine receptor CXCR3. Br J Pharmacol 166, 898–911 (2012).

49. Smith, J.S. et al. Biased agonists of the chemokine receptor CXCR3 differentially control chemotaxis and inflammation. Science Signaling 11, eaaq1075 (2018).

50. Stoffel, R.H., Randall, R.R., Premont, R.T., Lefkowitz, R.J. & Inglese, J. Palmitoylation of G protein-coupled receptor kinase, GRK6. Lipid modification diversity in the GRK family. J Biol Chem 269, 27791-4 (1994).

51. Premont, R.T., Inglese, J. & Lefkowitz, R.J. Protein kinases that phosphorylate activated G protein-coupled receptors. FASEB J 9, 175–82 (1995).

52. Lodowski, D.T., Pitcher, J.A., Capel, W.D., Lefkowitz, R.J. & Tesmer, J.J. Keeping G proteins at bay: a complex between G protein-coupled receptor kinase 2 and Gbetagamma. Science 300, 1256–62 (2003).

53. Jean-Alphonse, F. et al. Spatially restricted G protein-coupled receptor activity via divergent endocytic compartments. J Biol Chem 289, 3960–77 (2014).

54. Snyder, J.C. et al. Inhibiting clathrin-mediated endocytosis of the leucine-rich G protein-coupled receptor-5 diminishes cell fitness. J Biol Chem 292, 7208–7222 (2017).

55. Namkung, Y. et al. Monitoring G protein-coupled receptor and beta-arrestin trafficking in live cells using enhanced bystander BRET. Nat Commun 7, 12178 (2016).

56. Wright, S.C. et al. BRET-based effector membrane translocation assay monitors GPCR-promoted and endocytosis-mediated Gq activation at early endosomes. Proc Natl Acad Sci U S A 118(2021).

57. Koch, W.J., Inglese, J., Stone, W.C. & Lefkowitz, R.J. The binding site for the beta gamma subunits of heterotrimeric G proteins on the beta-adrenergic receptor kinase. J Biol Chem 268, 8256–60 (1993).

58. Gurevich, V.V. & Gurevich, E.V. GPCR Signaling Regulation: The Role of GRKs and Arrestins. Front Pharmacol 10, 125 (2019).

59. Touhara, K., Koch, W.J., Hawes, B.E. & Lefkowitz, R.J. Mutational analysis of the pleckstrin homology domain of the beta-adrenergic receptor kinase. Differential effects on G beta gamma and phosphatidylinositol 4,5-bisphosphate binding. J Biol Chem 270, 17000-5 (1995).

60. Zarca, A. et al. Differential Involvement of ACKR3 C-Tail in beta-Arrestin Recruitment, Trafficking and Internalization. Cells 10(2021).

61. Smit, M.J. et al. CXCR3-mediated chemotaxis of human T cells is regulated by a Gi- and phospholipase C-dependent pathway and not via activation of MEK/p44/p42 MAPK nor Akt/PI-3 kinase. Blood 102, 1959–65 (2003).

62. Grundmann, M. et al. Lack of beta-arrestin signaling in the absence of active G proteins. Nat Commun 9, 341 (2018).

63. Katada, T. & Ui, M. ADP ribosylation of the specific membrane protein of C6 cells by islet-activating protein associated with modification of adenylate cyclase activity. J Biol Chem 257, 7210–6 (1982).

64. Kurose, H., Katada, T., Amano, T. & Ui, M. Specific uncoupling by islet-activating protein, pertussis toxin, of negative signal transduction via alpha-adrenergic, cholinergic, and opiate receptors in neuroblastoma x glioma hybrid cells. J Biol Chem 258, 4870–5 (1983).

65. Hisano, Y. et al. Lysolipid receptor cross-talk regulates lymphatic endothelial junctions in lymph nodes. J Exp Med 216, 1582–1598 (2019).

66. Moller, T.C. et al. Dissecting the roles of GRK2 and GRK3 in mu-opioid receptor internalization and beta-arrestin2 recruitment using CRISPR/Cas9-edited HEK293 cells. Sci Rep 10, 17395 (2020).

67. Drube, J. et al. GPCR kinase knockout cells reveal the impact of individual GRKs on arrestin binding and GPCR regulation. Nat Commun 13, 540 (2022).

68. Ahn, S., Shenoy, S.K., Wei, H. & Lefkowitz, R.J. Differential kinetic and spatial patterns of beta-arrestin and G protein-mediated ERK activation by the angiotensin II receptor. J Biol Chem 279, 35518–25 (2004).

69. Zheng, H., Loh, H.H. & Law, P.Y. Beta-arrestin-dependent mu-opioid receptor-activated extracellular signal-regulated kinases (ERKs) Translocate to Nucleus in Contrast to G protein-dependent ERK activation. Mol Pharmacol 73, 178–90 (2008).

70. Harvey, C.D. et al. A genetically encoded fluorescent sensor of ERK activity. Proc Natl Acad Sci U S A 105, 19264–9 (2008).

71. Cheroutre, H. & Lambolez, F. Doubting the TCR coreceptor function of CD8alphaalpha. Immunity 28, 149–59 (2008).

72. Cole, D.K. et al. The molecular determinants of CD8 co-receptor function. Immunology 137, 139–48 (2012).

73. Thomsen, A.R.B. et al. GPCR-G Protein-beta-Arrestin Super-Complex Mediates Sustained G Protein Signaling. Cell 166, 907–919 (2016).

74. Nguyen, A.H. et al. Structure of an endosomal signaling GPCR-G protein-beta-arrestin megacomplex. Nat Struct Mol Biol 26, 1123–1131 (2019).

75. Cahill, T.J. et al. Distinct conformations of GPCR-β-arrestin complexes mediate desensitization, signaling, and endocytosis. Proc Natl Acad Sci U S^β^ A 114, 2562–2567 (2017).

76. Kavelaars, A. et al. Microglial GRK2: a novel regulator of transition from acute to chronic pain. Brain Behav Immun 25, 1055–60 (2011).

77. Singhmar, P. et al. Critical role for Epac1 in inflammatory pain controlled by GRK2-mediated phosphorylation of Epac1. Proc Natl Acad Sci U S A 113, 3036–41 (2016).

78. Kuai, J., Han, C. & Wei, W. Potential Regulatory Roles of GRK2 in Endothelial Cell Activity and Pathological Angiogenesis. Front Immunol 12, 698424 (2021).

79. Lymperopoulos, A., Rengo, G., Zincarelli, C., Soltys, S. & Koch, W.J. Modulation of adrenal catecholamine secretion by in vivo gene transfer and manipulation of G protein-coupled receptor kinase-2 activity. Mol Ther 16, 302–7 (2008).

80. Obrenovich, M.E. et al. Overexpression of GRK2 in Alzheimer disease and in a chronic hypoperfusion rat model is an early marker of brain mitochondrial lesions. Neurotox Res 10, 43–56 (2006).

81. Pitcher, J.A. et al. Role of beta gamma subunits of G proteins in targeting the beta-adrenergic receptor kinase to membrane-bound receptors. Science 257, 1264–7 (1992).

82. Pack, T.F., Orlen, M.I., Ray, C., Peterson, S.M. & Caron, M.G. The dopamine D2 receptor can directly recruit and activate GRK2 without G protein activation. J Biol Chem 293, 6161–6171 (2018).

83. Crilly, S.E. & Puthenveedu, M.A. Compartmentalized GPCR Signaling from Intracellular Membranes. J Membr Biol 254, 259–271 (2021).

84. Calebiro, D. & Godbole, A. Internalization of G-protein-coupled receptors: Implication in receptor function, physiology and diseases. Best Pract Res Clin Endocrinol Metab 32, 83–91 (2018).

85. Vilardaga, J.P., Jean-Alphonse, F.G. & Gardella, T.J. Endosomal generation of cAMP in GPCR signaling. Nat Chem Biol 10, 700–6 (2014).

86. Irannejad, R. & von Zastrow, M. GPCR signaling along the endocytic pathway. Curr Opin Cell Biol 27, 109–16 (2014).

87. Pavlos, N.J. & Friedman, P.A. GPCR Signaling and Trafficking: The Long and Short of It. Trends Endocrinol Metab 28, 213–226 (2017).

88. Mohammad Nezhady, M.A., Rivera, J.C. & Chemtob, S. Location Bias as Emerging Paradigm in GPCR Biology and Drug Discovery. iScience 23, 101643 (2020).

89. Thomsen, A.R.B., Jensen, D.D., Hicks, G.A. & Bunnett, N.W. Therapeutic Targeting of Endosomal G-Protein-Coupled Receptors. Trends Pharmacol Sci 39, 879–891 (2018).

90. Mai, Q.N. et al. A lipid-anchored neurokinin 1 receptor antagonist prolongs pain relief by a three-pronged mechanism of action targeting the receptor at the plasma membrane and in endosomes. J Biol Chem 296, 100345 (2021).

91. Jimenez-Vargas, N.N. et al. Agonist that activates the micro-opioid receptor in acidified microenvironments inhibits colitis pain without side effects. Gut 71, 695–704 (2022).

92. Latorre, R. et al. Sustained endosomal release of a neurokinin-1 receptor antagonist from nanostars provides long-lasting relief of chronic pain. Biomaterials 285, 121536 (2022).

93. Shukla, A.K., Xiao, K. & Lefkowitz, R.J. Emerging paradigms of beta-arrestin-dependent seven transmembrane receptor signaling. Trends Biochem Sci 36, 457–69 (2011).

94. Peterson, Y.K. & Luttrell, L.M. The Diverse Roles of Arrestin Scaffolds in G Protein-Coupled Receptor Signaling. Pharmacol Rev 69, 256–297 (2017).

95. Cahill, T.J 3rd., et al. Distinct conformations of GPCR-beta-arrestin complexes mediate desensitization, signaling, and endocytosis. Proc Natl Acad Sci U S A 114, 2562–2567 (2017).

96. Jiang, X., Yang, P. & Ma, L. Kinase activity-independent regulation of cyclin pathway by GRK2 is essential for zebrafish early development. Proc Natl Acad Sci U S A 106, 10183–8 (2009).

97. Sorriento, D. et al. The G-protein-coupled receptor kinase 5 inhibits NFkappaB transcriptional activity by inducing nuclear accumulation of IkappaB alpha. Proc Natl Acad Sci U S A 105, 17818–23 (2008).

