## Supplemental Figures for "Subcellular localization of GPCR kinases differentially modulate biased signaling at CXCR3"

SUPPLEMENTAL FIGURE 1

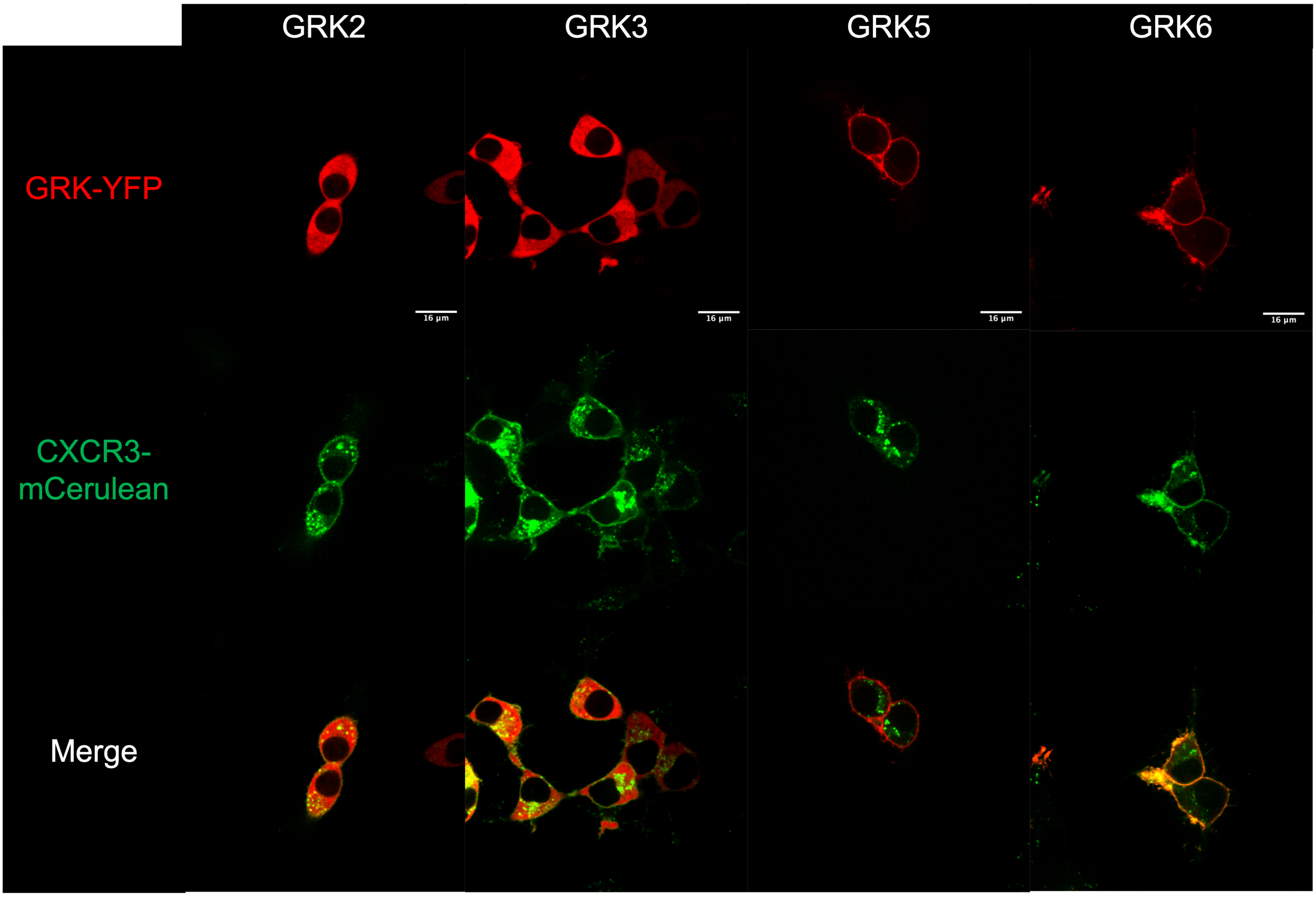

### SUPPLEMENTAL FIGURE 2

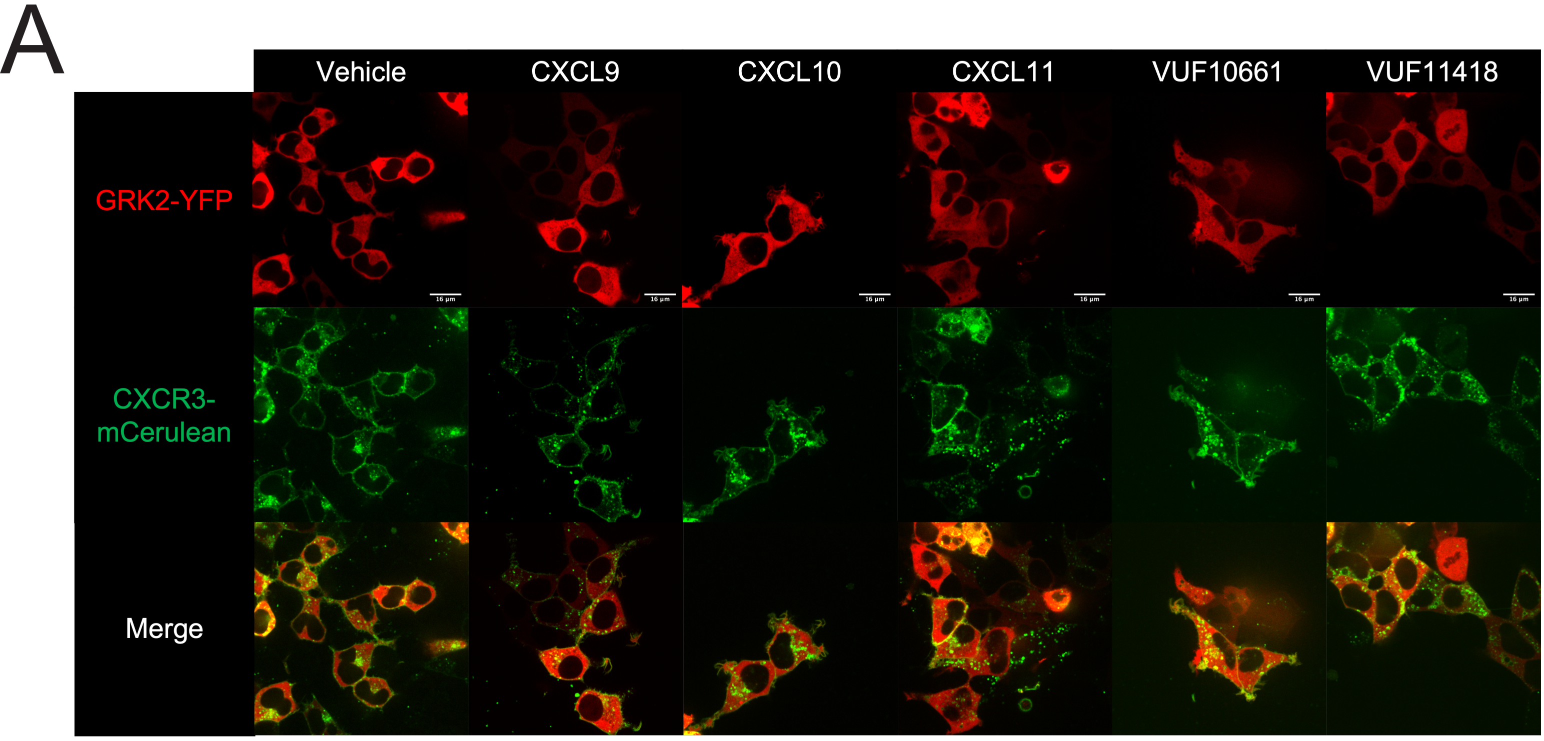


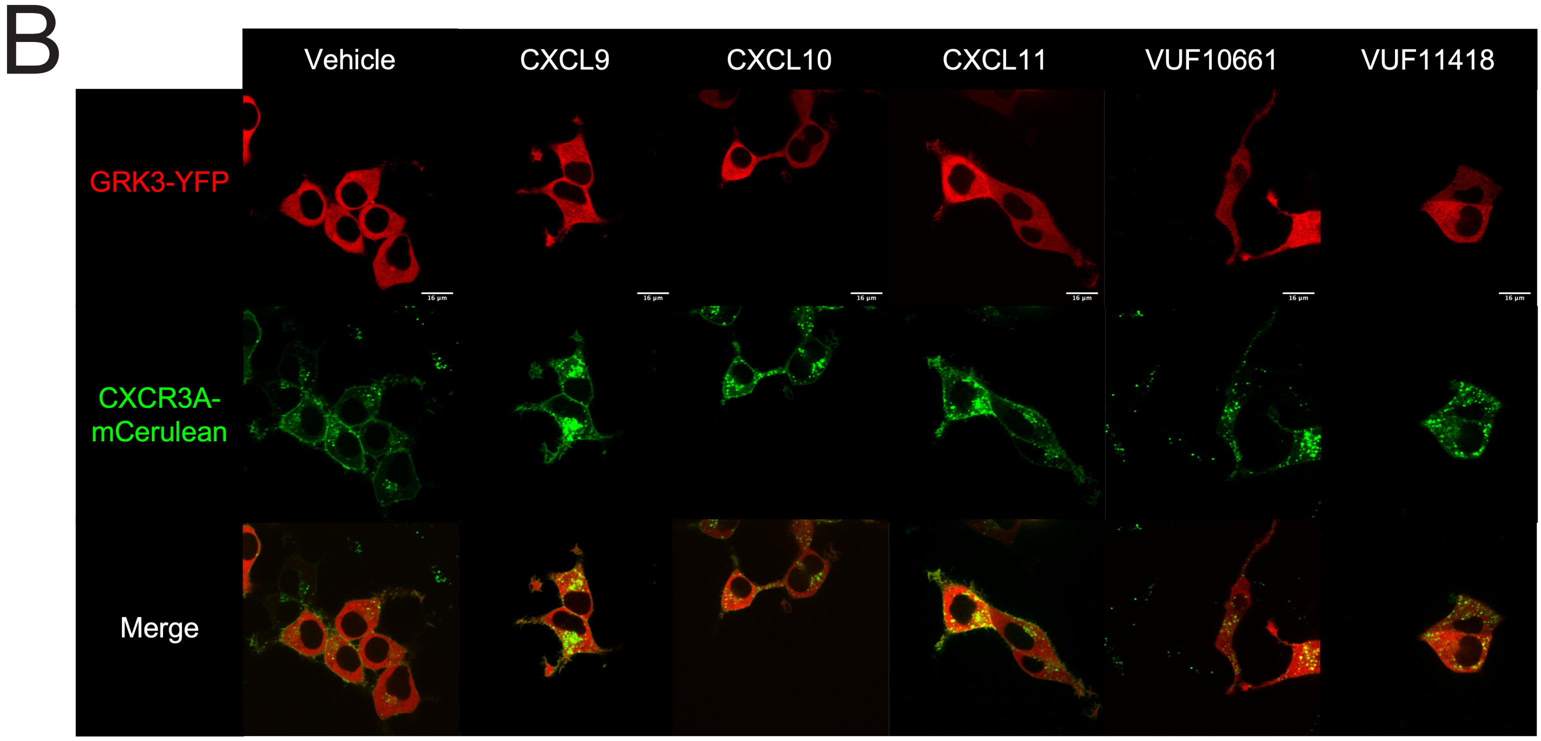

### SUPPLEMENTAL FIGURE 3

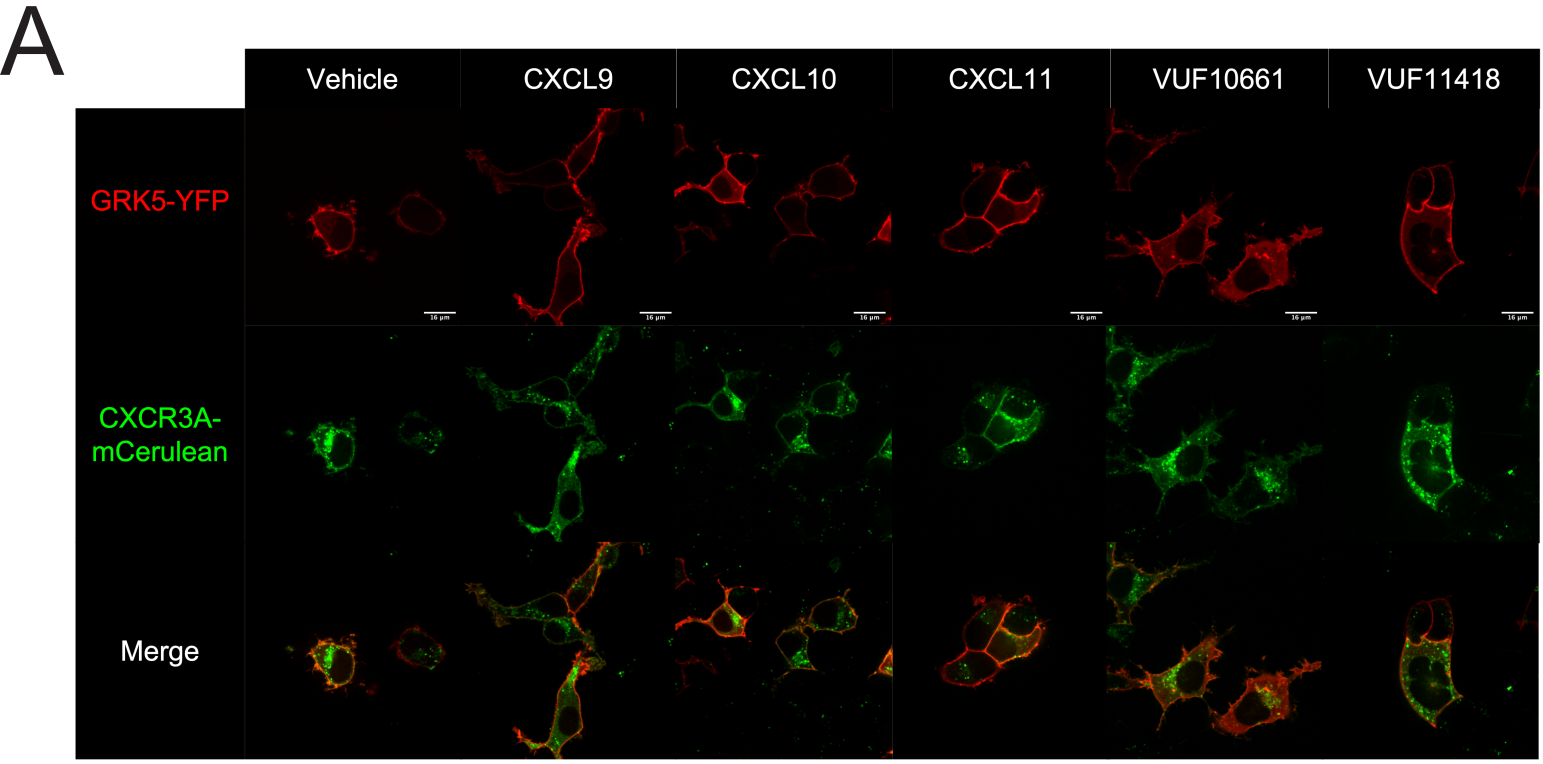


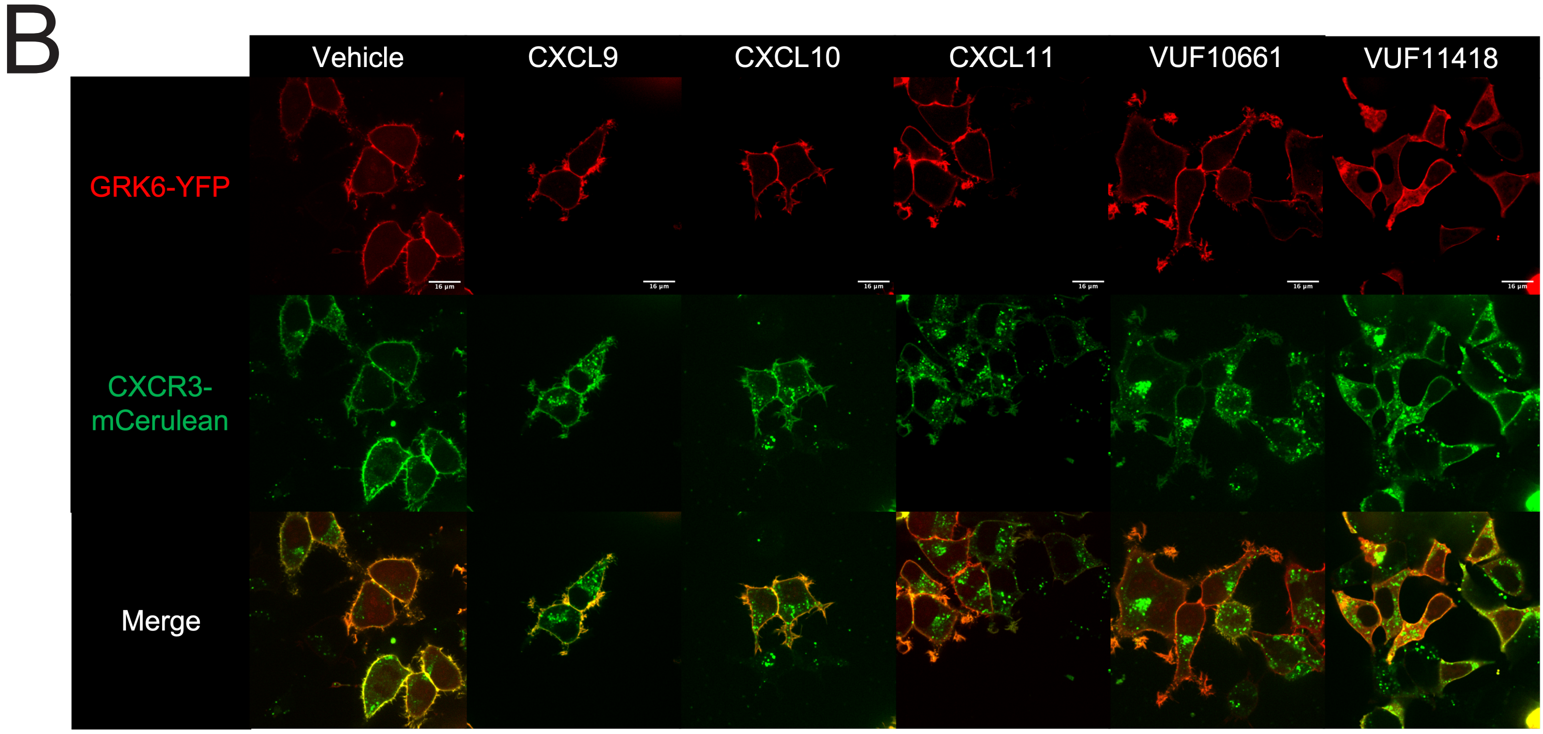

### SUPPLEMENTAL FIGURE 4

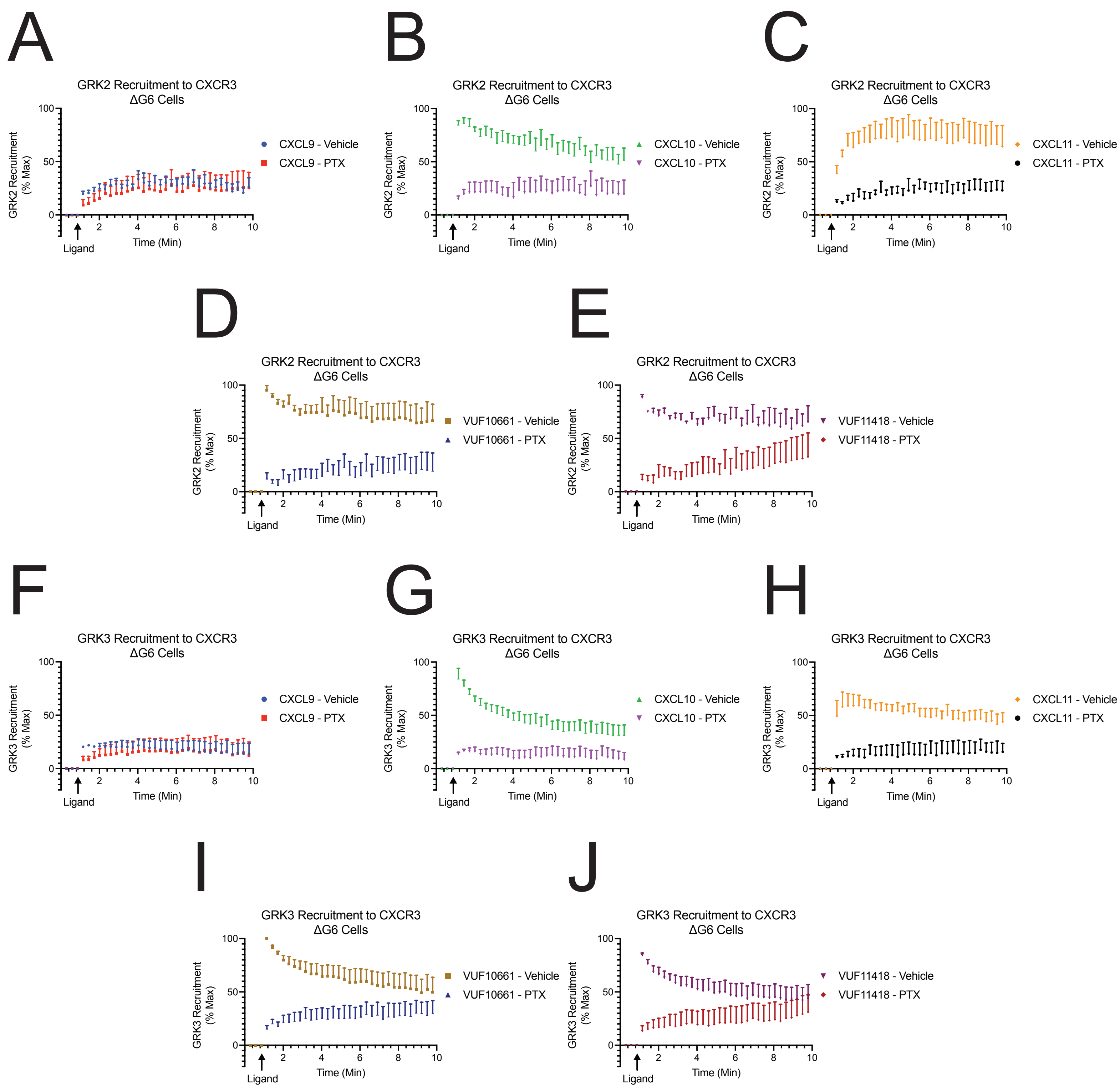

### SUPPLEMENTAL FIGURE 5

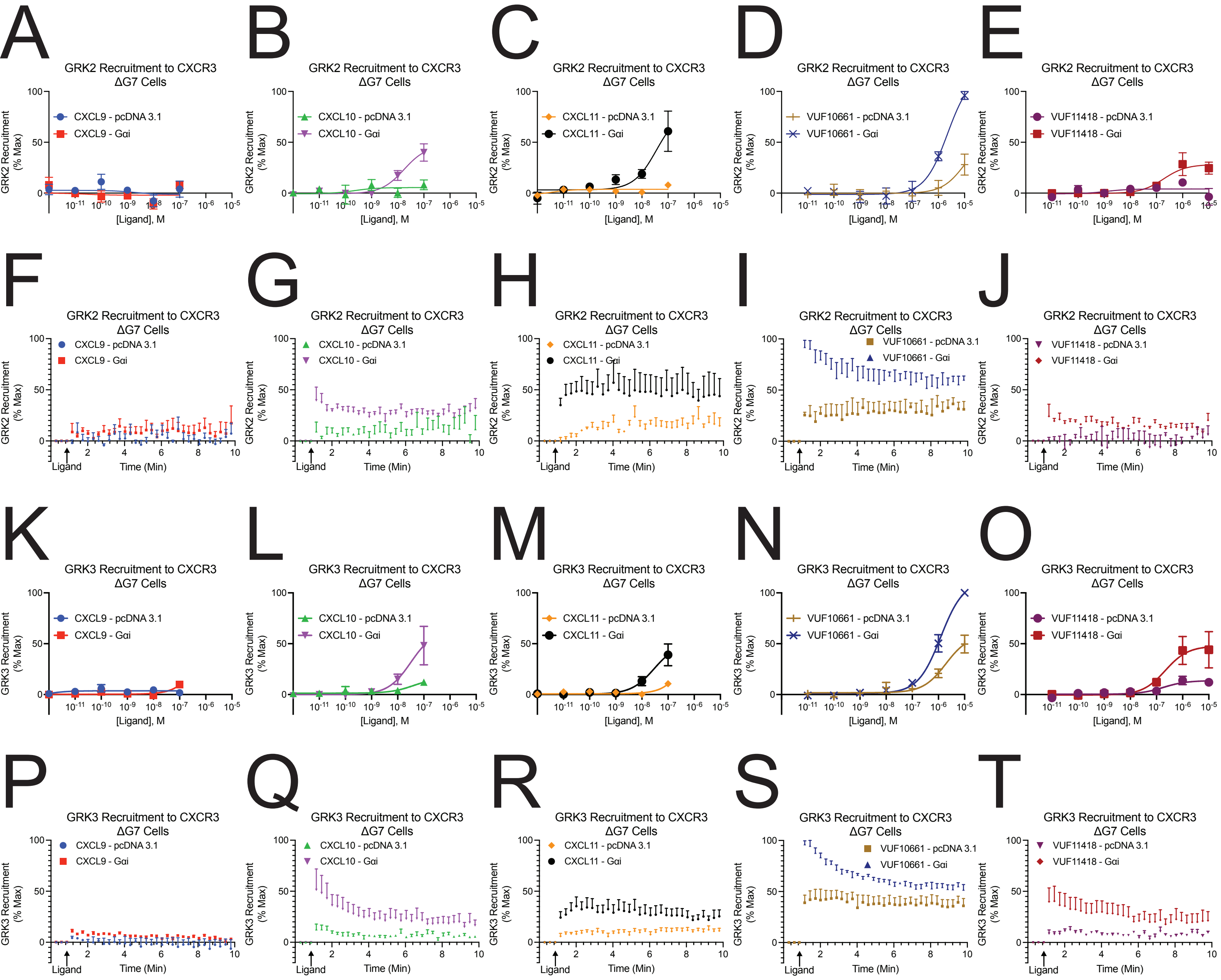

### SUPPLEMENTAL FIGURE 6

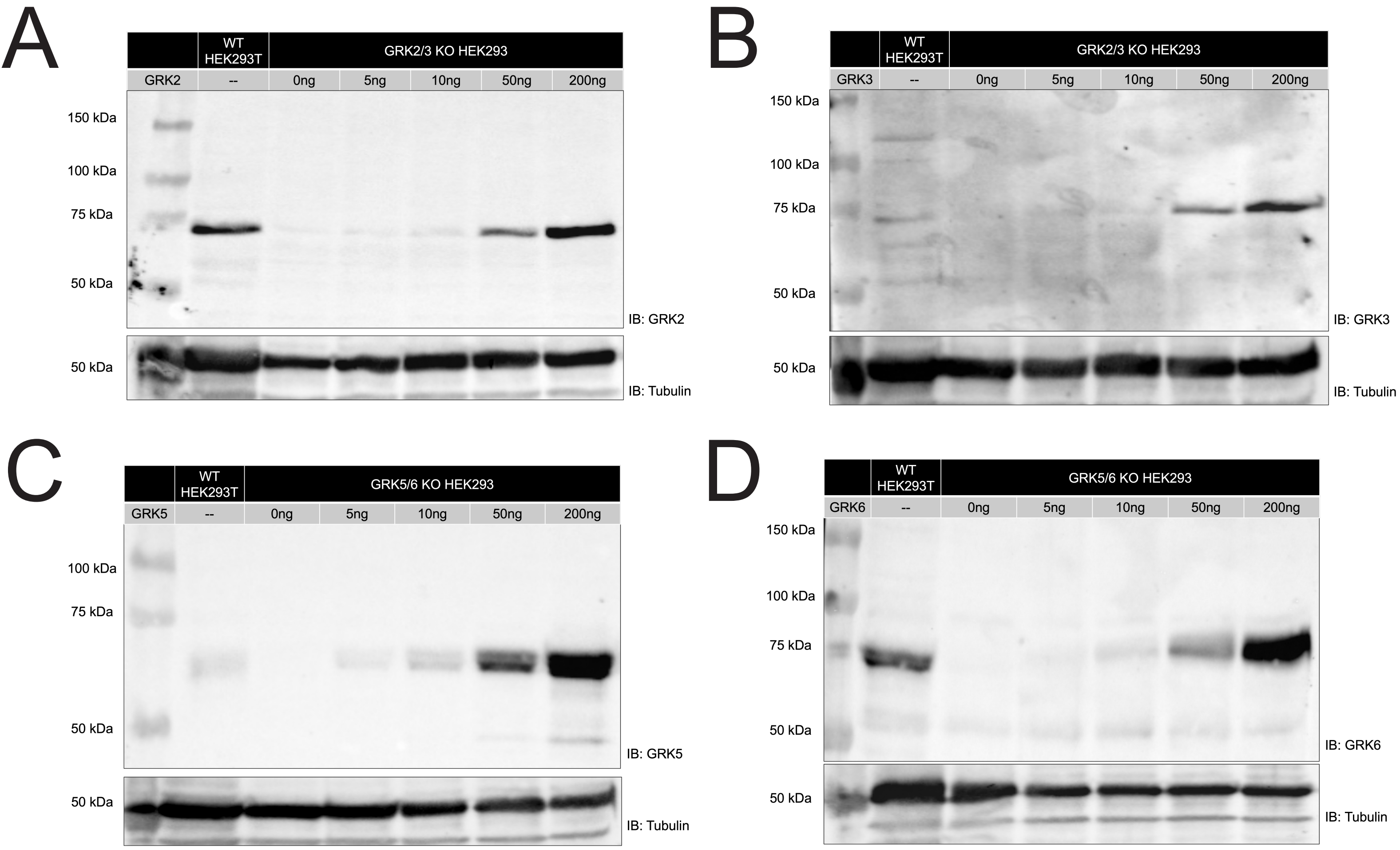

### SUPPLEMENTAL FIGURE 7

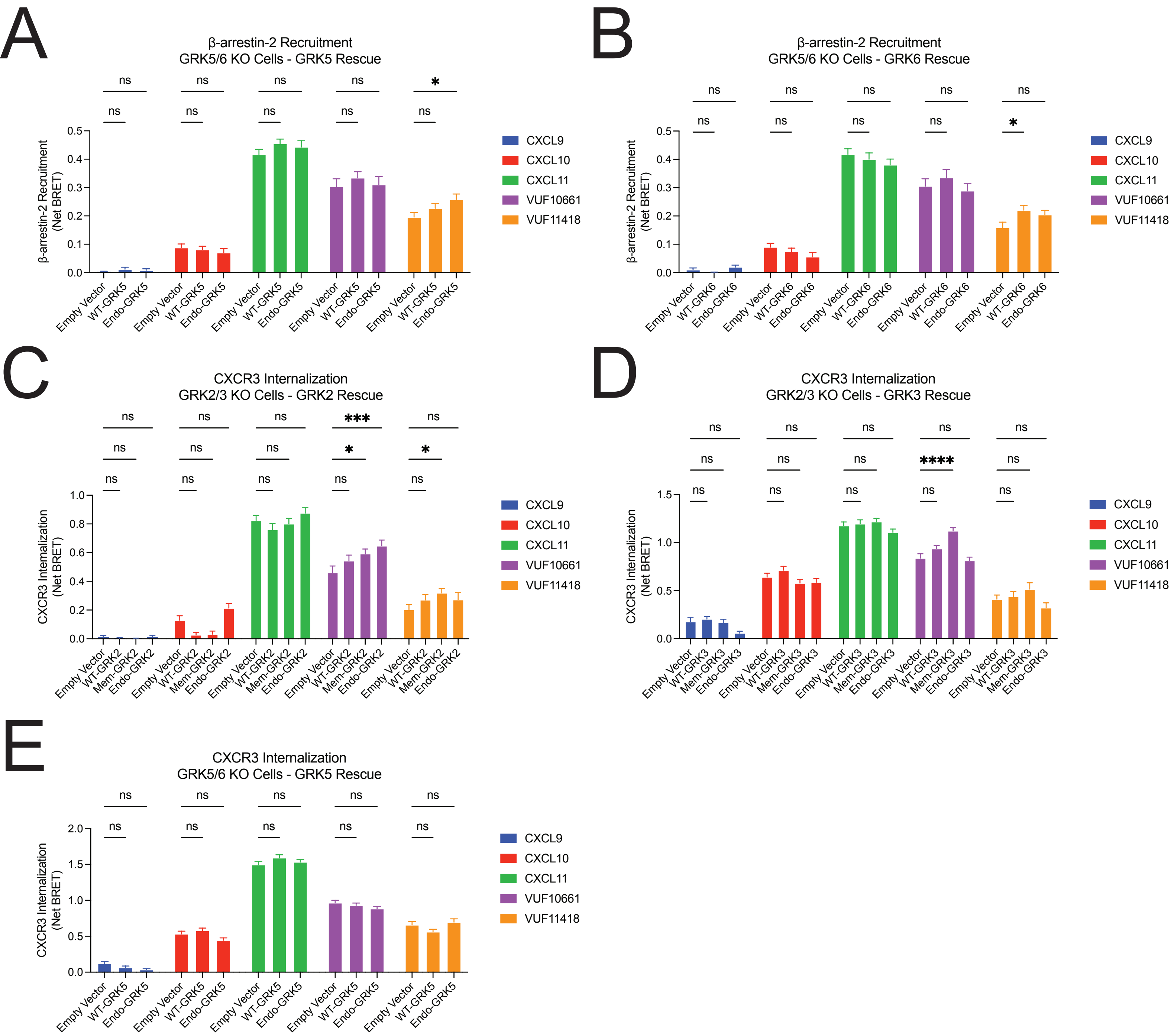
